# MODE: high-resolution digital dissociation with deep multimodal autoencoder

**DOI:** 10.1101/2025.01.02.631152

**Authors:** Jiao Sun, Ayesha A. Malik, Tong Lin, Ayla Bratton, Yue Pan, Kyle Smith, Arzu Onar-Thomas, Giles W. Robinson, Wei Zhang, Paul A. Northcott, Qian Li

## Abstract

In single cell biology, the complexity of tissues may hinder lineage cell mapping or tumor microenvironment decomposition, requiring digital dissociation of bulk tissues. Many deconvolution methods focus on transcriptomic assay, not easily applicable to other omics due to ambiguous cell markers and reference-to-target difference. Here, we present MODE, a multimodal autoencoder pipeline linking multi-dimensional features to jointly predict personalized multi-omic profiles and cellular compositions, using pseudo-bulk data constructed by internal non-transcriptomic reference and external scRNA-seq data. MODE was evaluated through rigorous simulation experiments and real multi-omic data from multiple tissue types, outperforming nine deconvolution pipelines with superior generalizability and fidelity.

## Background

Multimodal omics profiling and analysis, including transcriptomics, epigenomics, and proteomics, are crucial in cancer and immunology research. Initiatives like The Cancer Genome Atlas (TCGA) project and Clinical Proteomic Tumor Analysis Consortium (CPTAC) have performed bulk multi-omics analysis on various cancer types [1–3]. Recently, single-cell RNA sequencing (scRNA-seq) combined with other (epi)genomic data has enabled a deeper understanding of intra-tumoral heterogeneity, the identification of rare cell populations, and the exploration of tumor microenvironment interactions [4–7].

One challenge in single-cell analysis arises in lineage cell mapping for the developing human brains or Central Nervous System (CNS) tumors, such as pediatric medulloblastoma (MB) and other embryonal CNS tumors. Multiple studies have employed machine learning and deep learning algorithms to assign each cell in MB tumors to the closest matching cell state in a reference human embryonal cerebellum tissue [8, 9]. However, in Group 3 MB tumors, many cells were not successfully assigned to any predefined lineage cell state due to significant differences between the reference embryonal normal tissue and the target tumors. Moreover, this type of analysis may be more challenging for the tracing of regulatory or epigenomic landscapes (i.e., chromatin accessibility or DNA methylation [10]) in the origins of embryonal tumors because of the sparseness in single cell epigenomic data and the aforementioned problems in bioinformatics preprocessing.

Emerging technologies in single cell (multi-)omics like scATAC-seq, cyTOF, CITE-seq (protein and gene expressions) and single cell multiome (ATAC + gene expression) [11–14] offer promising avenues for the multi-dimensional integration at single cell resolution, enabling holistic understanding of complex biological processes. A well-known challenge in single cell multi-omics data preprocessing is the cell marker selection and cross-modality cell annotation. Tools like Seurat [15–18] and MaxFuse [19] have introduced innovative methods for cross-modality cell annotation in scATAC-seq or snmC-seq3 [20] based on reference scMultiome. These approaches may not be ideal for tumor tissues due to unexpected intra-tumoral and inter-tumoral heterogeneity.

An alternative approach for dissociating bulk multi-omics data and tracing lineage cells without single-cell profiling is to digitally deconvolute and purify bulk samples using an external single-cell reference. Most deconvolution methods [21–24] assume that the target bulk data and reference data share similar biological information at the cell state level, i.e., a signature matrix. If the target and reference datasets are generated from distinct platforms, normalization procedures can reduce discrepancies and enhance the accuracy of the output [25, 26]. A more advanced solution for addressing platform-related disparities is to train deep learning models, such as Scaden [27], TAPE [28], scpDeconv [29] on pseudo-bulk samples built from scRNA-seq reference, and then adapt the trained models to each target bulk sample. This type of normalization or adaptation in deconvolution may be insufficient to minimize profound reference-to-target disparities when tracing the lineage cell states for embryonal brain tumors, since the reference is derived from fetal normal brain tissues and the target profiles are tumors.

There are other challenges unresolved in these methods. First, to dissect epigenome or proteome, users must provide modality-matched single-cell references with preprocessed cell labels and inferred cell markers, such as single cell mass spectrometry or cyTOF proteomics [12, 30], scATAC-seq and bisulfite sequencing DNA methylation [31, 32]. However, existing deconvolution tools had not yet been benchmarked for dissecting bulk ATAC-seq data with scATAC-seq reference, whereas the pre-identified cell markers (peaks) may influence the accuracy of quantified cellular fractions. Second, the downstream differential (expression or accessibility) analysis between disease subgroups based on the digitally purified profiles (e.g., CIBERSORTx, TAPE) has not yet been rigorously validated by experiments. Lastly, most of deep learning deconvolution tools are trained based on pseudo-bulk data derived from external references, which may not adequately capture between-sample heterogeneity in the target bulk data. To improve the power and robustness of deconvolution, it is essential to incorporate the variability among target tissues into the training process.

To address the challenge of dissociating bulk epigenomes paired with transcriptomes, particularly when features (e.g., gene-peak or gene-CpG) are weakly linked, we propose the <u>M</u>ultim<u>O</u>dal <u>D</u>issociation via auto<u>E</u>ncoder (MODE) framework. MODE leverages cell markers detected in scRNA-seq data and accounts for the reference-to-target disparity, without requiring cell annotation across molecular modalities. To illustrate and benchmark the accuracy of MODE, we designed an extensive simulation study using two single cell multi-omic datasets from normal and tumor brain tissues [33, 34] along with the multimodal single-cell simulator scDesign3 [35–37], and then compared its performance with other tools: Scaden [27], TAPE [28], scpDeconv [29], DISSECT [38], BayesPrism [39], and CIBERSORTx [25], OmicVerse [40], scSemiProfiler [41]. For both tumor and normal bulk tissues, MODE consistently outperformed the competing methods, accurately quantifying cellular fractions in RNA and non-RNA molecules. We further assessed the accuracy of cell-type-specific differential expression (ctsDE) analysis by applying the Wilcoxon rank-sum test to in-silico transcriptomic and proteomic profiles of human cerebral cortex normal tissues digitally purified by competing methods. The results demonstrated that MODE yielded superior accuracy in downstream ctsDE detection by leveraging multimodal omics data. Finally, when applied to multiple brain tumor studies, MODE successfully uncovered different lineage cell states involved in the evolution of pediatric medulloblastoma (MB) tumor subgroups and benchmarked the prognostic value of glioblastoma multiforme (GBM) tumor microenvironment (TME) composition across molecular modalities. Our method was also applied to the TCGA breast cancer (BC) bulk ATAC-seq and DNAm data without using the tissue-paired bulk RNA-seq, illustrating its generalizability to the non-transcriptomic multimodal bulk sample deconvolution.

## Results

### Overview of MODE

We present MODE, a semi-reference based multimodal digital dissociation tool that resolves developmental origins or tumor microenvironment and recovers personalized multi-omic profiles mapping to each cell state (Figure 1a), generalizable to different molecules, tissue types, and diseases. In Step 1, MODE initialize the tissue-specific reference panels for the target bulk non-RNA profiles (i.e., epigenome or proteome) via the Joint Non-negative Matrix Factorization (JNMF) model and Projected Gradient Descent optimization [42], using the scaling factors (***s***_1_, ***s***_2_) to account for the modality-specific cell plasticity (Figure 1b). This step links the bulk RNA-seq and non-RNA profiles through tissue-specific cell counts fractions initialized by ***p***_0_ *~ Dir*(***π***, *v*) and then pre-purifies each bulk multiome, where the prior parameters (***π***, *v*) are the Dirichlet-Multinomial maximum likelihood estimate (MLE) on the external multi-subject cell counts with rows as subjects (or samples) and columns as annotated cell types.

**Fig. 1.**
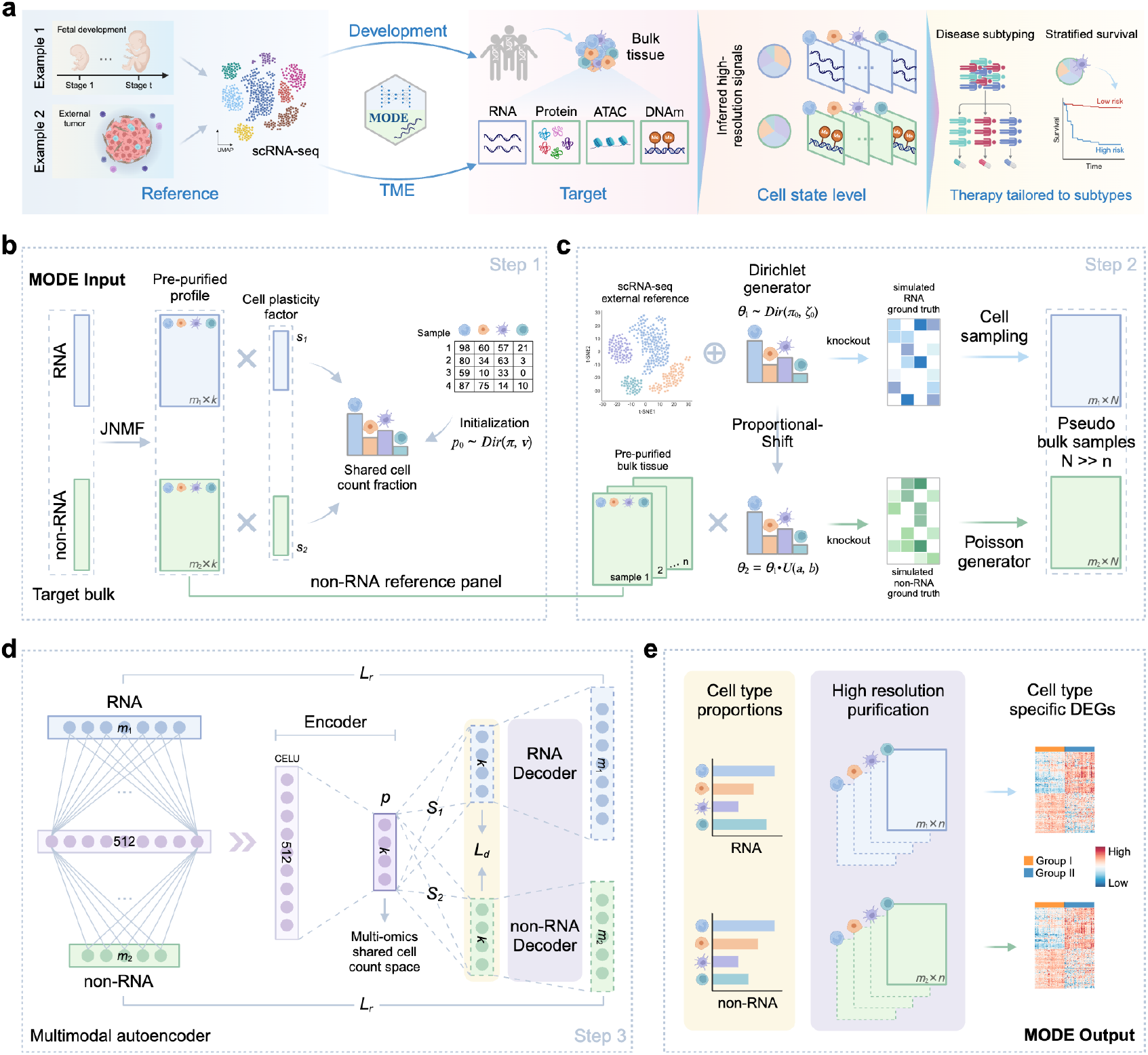
Overview of MODE framework. **(a)** MODE resolves the developmental origins or tumor microenvironment (TME) of target bulk tissues using scRNA-seq reference from fetal samples or external tumors. The dissociated multi-omic profiles may improve disease subtyping and prognosis prediction. **(b)** The input of MODE consists of RNA and non-RNA bulk omics data paired by tissues. MODE initializes a personalized and internal signature matrix via Joint Nonnegative Matrix Factorization and shared cell counts fractions. **(c)** Generate large-scale training data based on an external scRNA-seq reference and the internal signature matrices from Step 1. A proportional shift strategy coupled with modality-specific sparsity is used to simulate ground truth cellular compositions in RNA and non-RNA molecules. **(d)** A multimodal autoencoder with one encoder for the concatenated input data and two decoders for reconstructed bulk multiome. An updated personalized multi-omics signature matrix is extracted from two decoders adapted to each target tissue. **(e)** MODE outputs the predicted cell proportions and high-resolution purified multi-omic profiles, enabling downstream cts-differential analysis.

The second step of MODE (Figure 1c) aims to generate large-scale pseudo-bulk multiome data by connecting different molecular modalities through a proportion-shift strategy. Specifically, we first simulate the naive ground truth RNA cellular proportions without *a priori* information ***θ***_1_ *~ Dir*(***π***_0_, *ξ*_0_), where ***π***_0_ = (1, …, 1) and *ξ*_0_ is set at default value, similar to the approach in TAPE [28]. An external scRNA-seq reference with preprocessed cell labels is used to sample the cells per ground truth proportions, which are aggregated as a pseudo-bulk transcriptome. For the epigenome or proteome paired to transcriptome, the ground truth proportions ***θ***_2_ are generated by shifting RNA proportions ***θ***_1_, mimicking the shared cellular landscape and introducing cross-modality variability. The tissue-specific purified data recovered by JNMF in Step 1 are then used as internal reference panels to simulate the non-RNA pseudo-bulk profiles, with between-sample variability further enhanced by Poisson random generator. It should be noted that this data simulation procedure does not use the cell proportions estimated by JNMF in Step 1.

In Step 3 (Figure 1d), we employ a multimodal autoencoder for tissue-adaptive deconvolution, which integrates RNA and non-RNA bulk data to learn the shared cellular space. It consists of one encoder for the concatenated multi-omics input and two parallel decoders for reconstructing the input bulk data. The initial layer links the multimodal molecular features (***y***_1_ and ***y***_2_) through nonlinear CELU activation function:

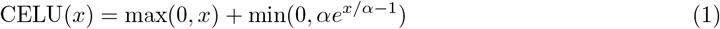

capturing the potential connections between cell marker transcription and cellular heterogeneity in other molecules. The encoder then further reduces the information to a low-dimensional embedding (***p***) representing the cell counts shared between molecular sources. To connect shared encoder with two decoders, this latent space representation is then linearly transformed into the modality-specific cellular composition and the reconstructed bulk multiome, via two separate fully connected layers ***S***_1_, ***S***_2_:

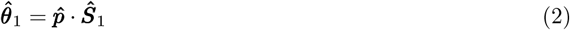

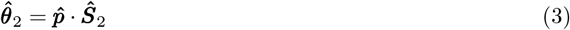

where 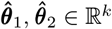 are estimated cell proportions of RNA and non-RNA; 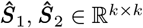 are the optimized weight matrices in fully connected layers and can be viewed as modality-specific cell size factors. Two decoders (*D*_1_, *D*_2_) are then leveraged to reconstruct the bulk data of each omics independently based on the cell proportions, each comprising five fully connected layers without activation function or bias. The decoders in molecular modality *i* (*i* = 1, 2) are determined by weight matrices 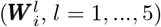 with an ReLU function to enforce non-negative values based on its biological meaning. 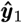 and 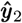 represent the reconstructed RNA and non-RNA bulk profiles with the same dimension as input data. 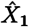 and 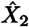 represent the predicted signature matrix of RNA and non-RNA, individually. That is

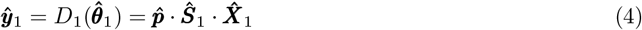

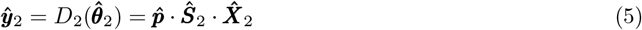

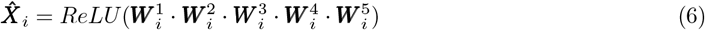

This framework is trained on the simulated bulk multi-omics data in Step 2 to minimize the loss in deconvoluted cellular composition and reconstructed bulk data for both modalities. In the adaptive stage, the parameters are refined for each target bulk sample to better capture the unseen patterns present in the tissue. It is worth noting that the feature selection in bulk epigenome or proteome does not require cell markers identified from non-RNA single cell reference. Instead, users can select the features from bulk data by feature-wise mean, variation, and sparseness. The output of MODE comprises modality-specific cellular compositions and high-resolution purified profiles (Figure 1e) that can be used in downstream differential analysis.

### Pseudo bulk multiome of developing human brain cerebral cortex

To benchmark our method, we first conducted an experiment on tissue-matched pseudo bulk multi-omics (gene expression and chromatin accessibility) profiles of developing human brain cerebral cortex, derived from real scMultiome data [33]. These brain tissues span six broad developmental periods (time points): early vs. late gestational fetal stages, infancy, childhood, adolescence, and adulthood. The scMultiome of these brain tissues is visualized in Figure 2a, showing high-quality cell annotation across modalities, such as excitatory neuron (EN), inhibitory neuron (IN), oligodendrocyte progenitor cell (OPC), and oligodendrocyte. More than 99% of cells in Seurat clusters EN-fetal-early, EN-fetal-late, IN-fetal are from fetal samples, while clusters EN and IN were exclusively from the non-fetal samples. Hence, it is reasonable to assume the rareness or absence of embryonal neuron cells in the neocortex of children, adolescents, and adults.

**Fig. 2.**
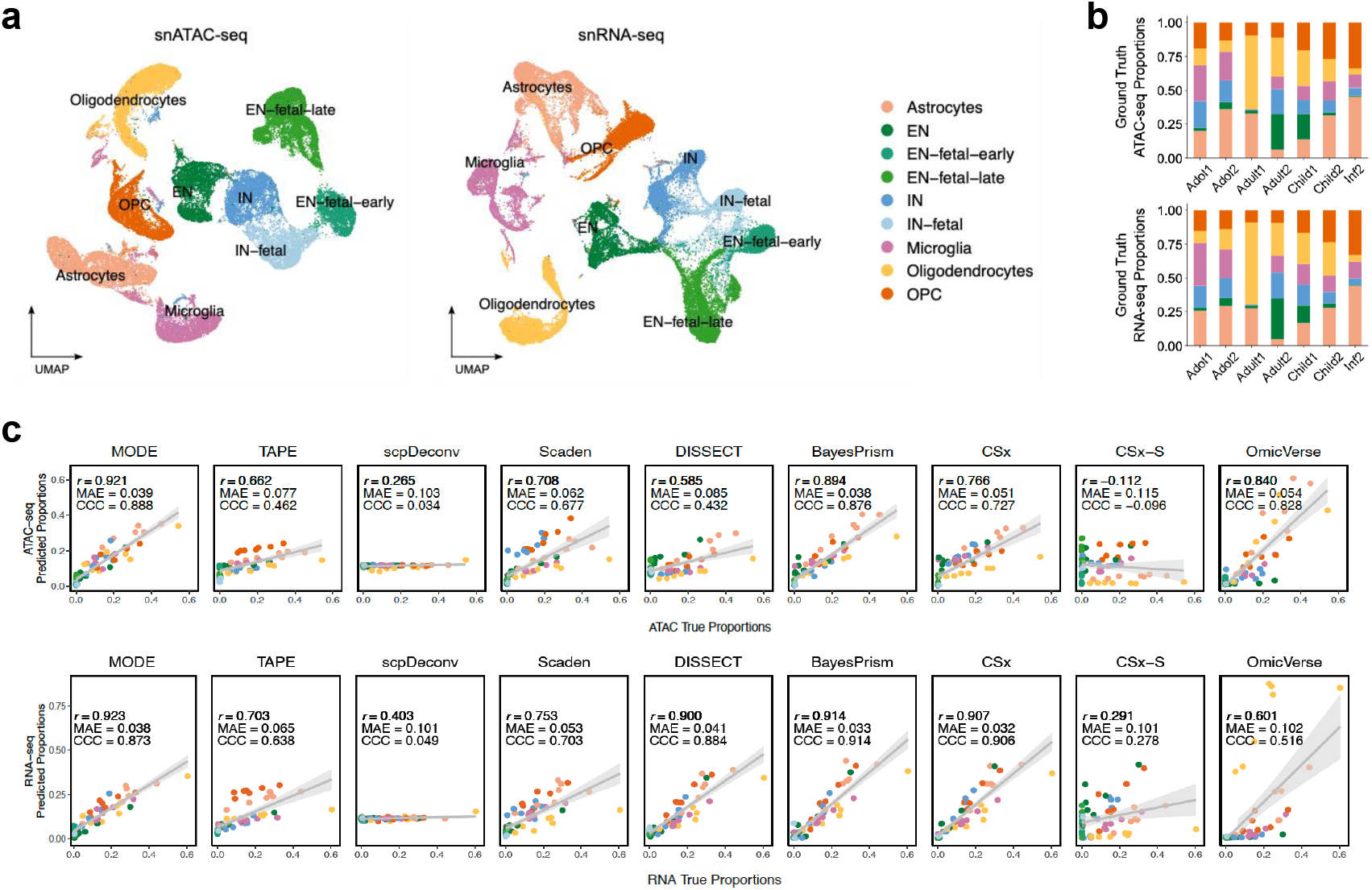
Deconvolution of pseudo bulk multiome in human brain cerebral cortex. **(a)** Visualization of chromatin accessibility and gene expression paired at single cell resolution in twelve developing human neocortex samples by Uniform Manifold Approximation and Projection (UMAP). **(b)** Bar plots for ground truth cell proportions in each pseudo bulk sample. **(c)** Scatter plots for cell proportions estimated by MODE and competing methods vs. ground truth, assessed by Lin’s CCC, MAE and Pearson correlation. EN: excitatory neuron; IN: inhibitory neuron; OPC: oligodendrocyte progenitor cell.

We constructed *n* = 7 postnatal pseudo bulk samples at the stages of infant, child, adolescent, and adult as target data, with ground truth cell fractions visualized in Figure 2b. For each sample, the simulated cell proportions in transcriptome vs. epigenome are different. Next, we used external scRNA-seq and scATAC-seq profiles at embryonal (*n* = 2) and infant (*n* = 1) time points as deconvolution reference basis. For each molecular modality, we selected the cell-type-specific marker genes or peaks from the reference single-cell profiles using differential analysis in Seurat [17] and then subset the reference and target bulk data to the pre-identified cell markers.

To illustrate the power and robustness of MODE, we thoroughly permuted 1296 different scenarios of hyperparameters, evaluating deconvolution accuracy by Lin’s Concordance Correlation Coefficient (CCC), Mean Absolute Error (MAE), and Pearson correlation (r). We also used multi-subject cell counts derived from all vs. postnatal samples to initialize the JNMF in Step 1. The results in Supplementary File 1 implied that MODE yielded high accuracy in deconvoluted cell fractions regardless of hyperparameters when the input cell counts for Dirichlet initialization (in JNMF) were extracted from reference samples similar to the target (non-fetal) tissues. The JNMF purified ATAC data was compared to the ground truth CTS profiles using Pearson correlation (Supplementary File 1 and Supplementary Figure S1), illustrating that JNMF partially recovered the biological information in the target tissues. Next, we compare MODE with 9 deconvolution methods: CIBERSORTx (CSx), CSx-S mode, TAPE, Scaden, BayesPrism, scpDeconv, DISSECT, OmicVerse, and scSemiProfiler in each modality. Notably, MODE is one of the top performers in both RNA-seq and ATAC-seq based on Pearson correlation (*r* = 0.923, 0.921, respectively), while BayesPrism (CCC = 0.914, MAE = 0.033) marginally outperforms MODE (CCC = 0.873, MAE = 0.038) in RNA-seq deconvolution. The cellular proportions resolved by CSx (without normalization) and DISSECT (i.e., a variational autoencoder framework) are desirable in RNA-seq (CCC = 0.906, 0.884; *r* = 0.907, 0.900) and acceptable in ATAC-seq (CCC = 0.727, 0.432; *r* = 0.766, 0.585). The other two deep learning-based methods, Scaden and TAPE, also predicted cellular fractions with acceptable performance in at least one modality (CCC = 0.703, 0.638 in RNA-seq, 0.677, 0.462 in ATAC-seq; *r* = 0.753, 0.703 in RNA-seq, 0.708, 0.662 in ATAC-seq). On the other hand, the normalization procedure implemented in CSx-S mode failed to improve the cell fractions in both RNA-seq and ATAC-seq due to overfitting. The tool scpDeconv designed for proteomics deconvolution did not yield acceptable result in each modality (CCC = 0.049, 0.034), while OmicVerse performed well in ATAC-seq (*r*=0.840, CCC=0.828) but not in RNA-seq (*r*=0.601, CCC=0.516) at its default configuration. The last competing method scSemiProfiler digitally dissociated the bulk samples at single cell level and required a subset of target tissues profiled by both bulk and single cell assays. This tool is not directly comparable to the other competing methods because it requires tissue-paired bulk and single-cell data and downstream clustering of dissociated cells. The results of scSemiProfiler were shown in Supplementary Figures S2-S3.

Another objective of this experiment is to determine a range of default configurations for our tool. The results in Supplementary File 1 suggest that MODE yields optimal overall performance at subj var=0.05 and batch size=64 for this data, regardless of the other hyperparameters. The parameter subj var represents the variation in cellular compositions between target tissues, that is, a larger value accounts for more heterogeneity. Therefore, a tentative default configuration of MODE Python package is subj var=0.05, batch size=64, epochs=250 for non-tumor deconvolution, which is subject to fine-tuning for tumor tissues. It should be noted that we ran each baseline method using the default configuration of each tool, while the MODE result shown in Figure 2c is the best among the permuted hyperparameters (Supplementary File 1). Hence, the results of some baseline tools may be improved by fine-tuning. The superior performance of MODE, BayesPrism, and OmicVerse in ATAC-seq should be further investigated in tumor tissues, whereas the single cell annotation and cell marker identification in reference data are challenging and prone to errors.

### In-silico gene expression and chromatin accessibility in glioblastoma tumors

To evaluate each deconvolution method in tumor tissues with sufficient sample size, we designed a novel simulation study using scDesign3 [37], visualized in Supplementary Figure S4. This pipeline generates gene expression and chromatin accessibility paired by in-silico single cells, which are aggregated to construct pseudo bulk data per cell state. To mimic the variability among tumor tissues, we utilized the real scRNA-seq and scATAC-seq data paired by GBM tumor samples [34] to estimate the statistical parameters required by scDesign3, visualized in Supplementary Figure S4. The cell labels of this real scATAC-seq data were predicted by Seurat [17], which transferred the scRNA-seq cell labels to scATAC-seq in the same biological system. Although this multimodal cell annotation algorithm cannot label some GBM malignant cells precisely in scATAC-seq according to paired UMAPs (Supplementary Figure S4a), it does not affect the overall statistical parameters extracted from gene expression and chromatin accessibility per cell type in the next step (Supplementary Figure S4b). The synthetic single cells per modality generated from scDesign3 are shown in Supplementary Figure S4c. The mean local inverse Simpson’s index (mLISI) measuring the similarity between real and synthetic cells were shown in the UMAP (Supplementary Figure S4c), whereas mLISI=1.89 in scRNA-seq and mLISI=1.64 in scATAC-seq. The details of this simulation process are described in Methods.

We generated pseudo bulk target samples (n=100) of small (Supplementary Figure S5a) vs. large (Supplementary Figure S5b) difference between modalities, with details explained in Methods. It should be noted that this experiment aims to assess the accuracy of each deconvolution method without epigenomic cell markers pre-identified from scATAC-seq reference. Hence, the scATAC-seq features (peaks) detected in *>* 10% cells were all selected for epigenomic deconvolution regardless of cell marker identification, eliminating the sparseness in scATAC-seq reference. To demonstrate the validity of this simulation design, the cell marker genes used in transcriptomic deconvolution were still selected from reference scRNA-seq data [34] using differential analysis in Seurat.

A hyperparmeter was slightly tuned (i.e., subj var=0.1) in this experiment and the other three studies of tumor deconvolution because of the well-known intra- and inter-tumoral heterogeneity. For each target bulk data, MODE can accurately estimate RNA-seq and ATAC-seq cell proportions (Figure 3a-b). Most of the competing methods can deconvolve bulk transcriptome with scRNA-seq reference and cell markers. Specifically, BayesPrism, CSx, CSx-S mode, DISSECT, TAPE, and Scaden achieved better performance in RNA-seq deconvolution (Lin’s CCC *>* 0.9) compared to the previous experiment due to the lower disparity between reference and target in the current experiment. However, these competitors failed to deconvolve pseudo bulk ATAC-seq data (Supplementary Figure S5) due to the lack of cell markers and/or sparseness in reference scATAC-seq. Surprisingly, scpDeconv - a tool having poor performance in the experiment of developmental human brains - was the second-best performer in deconvolving GBM tumor multiome with CCC = 0.587, 0.432 for ATAC-seq and CCC = 0.481, 0.540 for RNA-seq, respectively (Supplementary Figure S5). The difference of scpDeconv performance between two experiments can be explained by the increase of target data sample size and the algorithm implemented in this tool imputing single cell reference before deconvolution, which favors reference data with less sparseness.

**Fig. 3.**
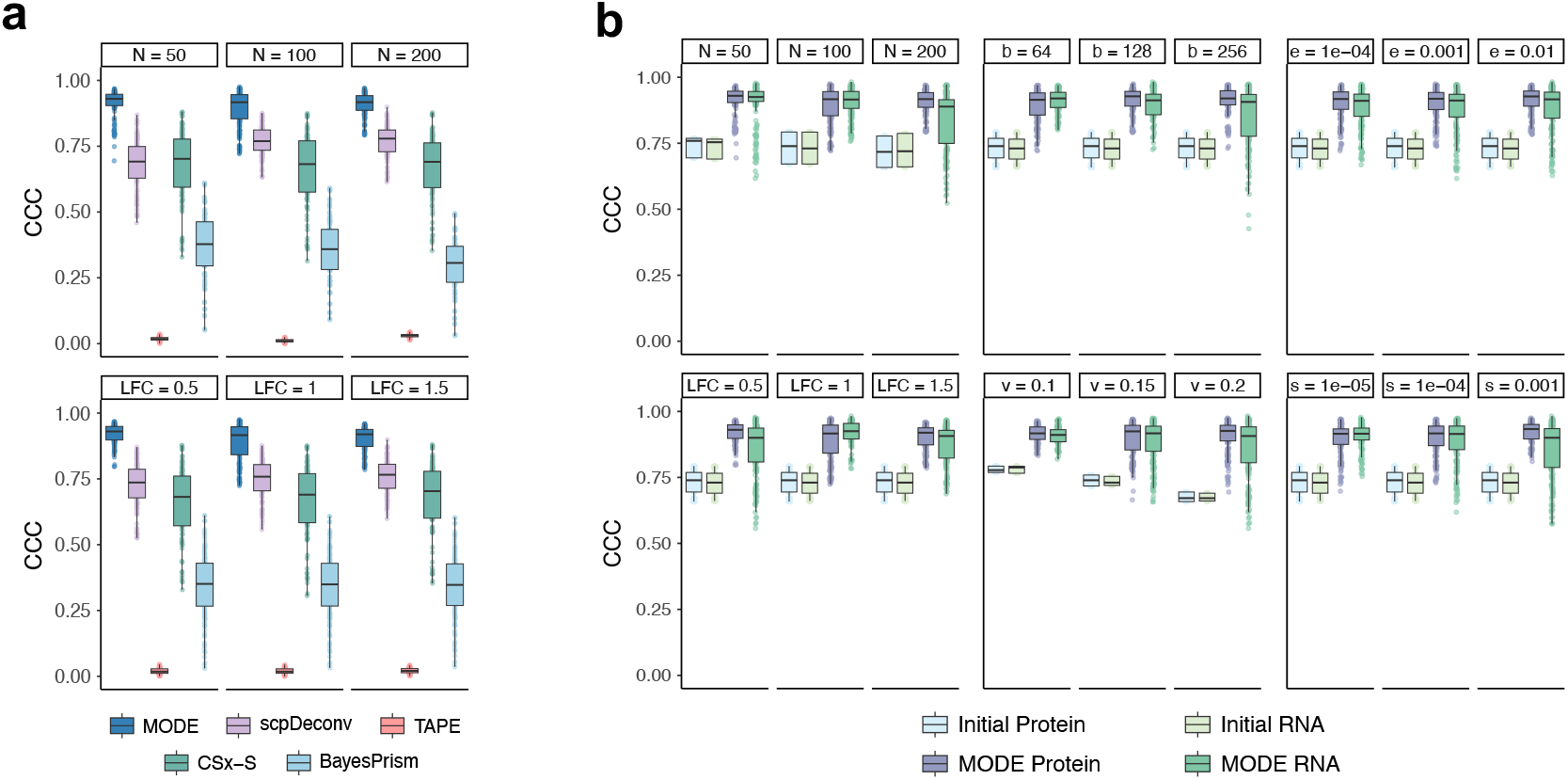
Performance on simulated proteomics data from human brain normal tissues. **(a)** CCC of MODE, scpDeconv, TAPE, CSx-S, and BayesPrism on bulk proteomics deconvolution. **(b)** CCC of MODE and the initial cell count fractions in Step 1 JNMF purification by different normal tissue simulation scenarios: sample size (N), log fold change (LFC); MODE hyperparameters: batch size (b), initial proportion dispersion (v), step size (s), and convergence threshold (e).

### In-silico bulk transcriptomics and proteomics

The above simulation studies only used pseudo bulk transcriptome and epigenome, without generating real bulk RNA-seq or other omics data, such as mass spectrometry (MS) proteomics. Another critical component in MODE is the high-resolution purification of multi-omics, i.e., predicting cell-type-specific profiles for each tissue. Therefore, to examine deconvolution in other omics and benchmark the quality of purification, we designed another experiment to generate nine scenarios of bulk RNA-seq transcriptome and MS proteome from adult human brain prefrontal cortex normal tissues in multiple sclerosis, using snRNA-seq data [43] along with pure-cell and mixed-cell proteomes [44, 45]. Each scenario consisted of a target bulk multi-omic dataset defined by a sample size *N* = 50, 100, 200 and cts-differential expression (ctsDE) log fold change (LFC = 0.5, 1, 1.5) between diseased and controls. Details of this simulation design were described in previous work [42].

We compared MODE with deconvolution methods that impute or adapt the reference according to target bulk data: scpDeconv, TAPE, and CSx-S mode, BayesPrism (Figure 3a and Supplementary Figure S6), using *N* = 50 single cell references randomly selected from the same scRNA-seq data [43] to rerun each tool 50 times on the target bulk samples. Further, we also reran MODE repeatedly for each scenario by permuting different hyperparameters, such as between-subject variance (v) in initial cell fraction, step size (s) and convergence threshold (e) in Step 1, and batch size (b) in Step 3. TAPE had poor performance with average CCC = 0.020 across all scenarios, revealing that this method cannot handle the cross-modality difference between reference and target bulk profiles. The imputation and domain-adversarial training in scpDeconv and the cross-platform normalization in CSx-S narrowed the gap between reference scRNA-seq and target bulk proteomics data with acceptable accuracy in cell abundance quantification (i.e., mean CCC = 0.738, 0.644), although the variation of CSx-S performance was very large. MODE consistently outperformed TAPE, scpDeconv, CSx-S, and BayesPrism in cell abundance quantification for bulk proteomics across all scenarios (Figure 3a and Supplementary Figure S6). In addition, MODE has yielded superior performance in cell abundance quantification for both omics across all scenarios and hyperparameters, substantially improving the accuracy compared to the initial cell counts fractions in terms of CCC and MAE (Figure 3b and Supplementary Figure S7). Overall, MODE demonstrated stable cell fraction estimates regardless of the variation in initial values. Our multimodal autoencoder remained powerful for a wide range of hyperparameters used in the JNMF initialization and parallel decoder training.

In the assessment of high-resolution purification, we only consider MODE, TAPE, BayesPrism, and CSx-S since scpDeconv does not purify bulk profiles. The evaluation metrics are described in Methods. The dimensions of multimodal expression profiles purified by CSx-S were extremely low with 58% of datasets yielding *<* 300 valid features, which cannot be used in downstream evaluation (Supplementary File 3). Among the high-resolution multiomes purified by MODE with different hyperparameters, we filtered out the results with *<* 300 valid features. For each method, the top ten F1-score for ctsDEG and ctsDEP in purified transcriptome and proteome were visualized in Figure 4a-b, demonstrating overall superior performance of MODE across different sample sizes and LFCs, with BayesPrism leading in some scenarios. Specifically, as sample size or LFC increases, the Wilcoxon test on MODE-purified profiles yields better F1-score for neuron in both protein and RNA expressions, while BayesPrism achieves higher scores for both protein and RNA expressions in astrocytes. The accuracy of ctsDEG in TAPE-purified RNA-seq profiles does not increase at larger sample size and LFC.

**Fig. 4.**
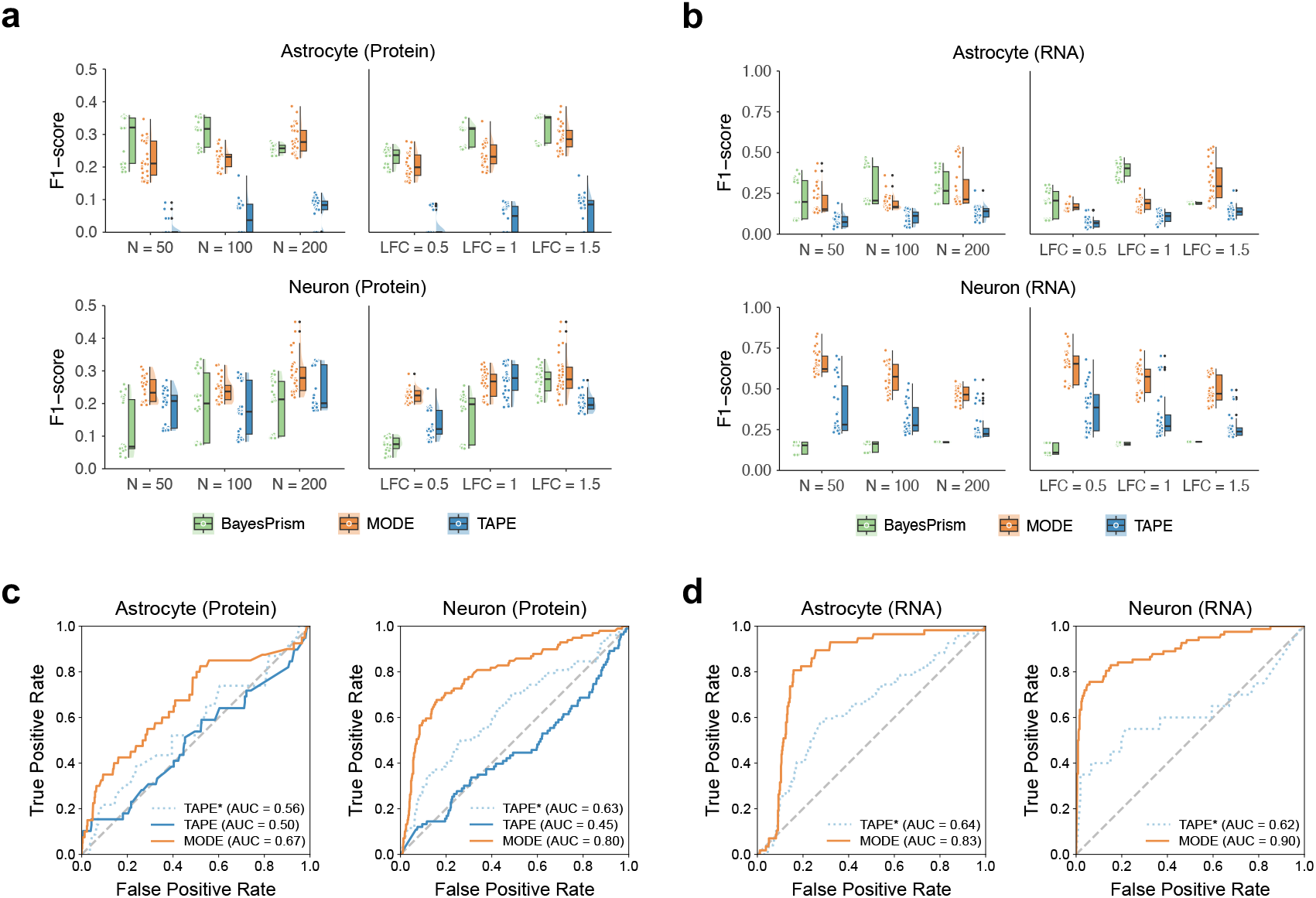
Cell-type-specific multiome profiles recover differentially expressed features. **(a, b)** Raincloud plots for F1-score on ctsDE detection in Astrocyte- and Neuron-specific proteome and transcriptome by MODE, TAPE, and BayesPrism. **(c, d)** Area Under the Receiver Operating Characteristic Curve (AUC) for the best F1-score of each method based on the same target data (N=200, LFC=1.5). Solid lines represent the autoencoder hyperparameter setting used in the best result of MODE, which may or may not be identical to the default setting used in TAPE (dotted line).

The results in Figure 4a demonstrated that MODE was more robust and powerful in predicting protein and RNA expression in neurons compared to BayesPrism and TAPE. The neuron-specific RNA-seq gene expression recovered by MODE, TAPE, BayesPrism yielded higher true positive rate (TPR) compared to the astrocyte-specific transcriptomics or proteomics, although the false positive rate (FPR) in the scenarios of large sample size or LFC was relatively higher with lower F1 score (Figure 4b). Moreover, the poor F-score (*<* 0.3) in most of the results purified by MODE, TAPE, BayesPrism except MODE neuron-specific RNA, and BayesPrism astrocyte-specific protein (Figure 4a-b), was primarily driven by the inflated FPR (*>* 0.4), although TPR of MODE was ideal and better than TAPE in some scenarios, as shown in Supplementary File 3. Meanwhile, we selected the results of MODE and TAPE from similar or identical hyperparameters for the scenario of maximum sample size (N=200) and LFC=1.5. The Area Under the Curve (AUC) illustrated significant improvement by MODE on DE detection in astrocyte and neuron cells across modalities, visualized in Figure 4c-d. In general, MODE substantially improved the quality of high-resolution ctsMultiome by leveraging non-RNA profiles in the target tissues.

### Pediatric medulloblastoma RNA-seq and DNAm

Medulloblastoma is one of the most malignant childhood brain tumors with diversified treatment responses and stratified outcomes. A recent study [8] mapped pediatric MB tumors to the human fetal cerebellum origins by deconvolving the bulk transcriptome of tumor tissues collected in the International Cancer Genome Consortium (ICGC) [46] and the SJMB12 cohort. SJMB12 is a multi-center clinical trial enrolling 660 children ages 4-25 with newly diagnosed medulloblastoma. Importantly, this trial was the first to incorporate molecular subgrouping into risk stratification, tailoring therapy to both biological and clinical features. The deconvolution tool MuSiC [23] and a scRNA-seq reference data from human fetal cerebellum normal tissues [47] were deployed to deconvolve bulk RNA-seq data from these cohorts [8]. The cellular compositions in transcriptome was found to correlate with MB subgroups clinically classified based on DNAm signatures: Sonic Hedgehog (SHH), Group 3 (G3), and Group 4 (G4). However, cell types such as Astro/Glia and GlutaCN/UBC—identified as progenitors or the lineage of origin for G3 and G4 MB malignant cells in scRNA-seq analyses [48]—were not fully recovered by MuSiC in the bulk transcriptome due to computational limitations.

The paired methylome-transcriptome data from both cohorts have been previously profiled with detailed preprocessing descriptions [8, 49]. Hence, we applied MODE to the bulk multi-omics profiles of n=253 MB tumors, using the above human developmental cerebellum scRNA-seq data as reference. We used 3146 cell type marker genes identified from this reference. From the 10,000 most variable CpGs in the DNAm data, we first selected those linked to the cell marker genes using CpG-gene mapping in MethylMix [50] and then further subset to 1000 CpGs contributing to the latent cellular embeddings shared across modalities using AJIVE algorithm implemented in MICSQTL [42]. We controlled the dimension of DNAm in this analysis to optimize the predicted origin-specific profiles. The deconvoluted cell fractions and purified gene expression were compared between MB subgroups using the Wilcoxon rank-sum test and BH correction for FDR.

The human fetal cerebellum derived cell states mapping to MB tumors were quantified by MODE in both methylome (Figure 5a) and transcriptome (Supplementary Figure S8), outperforming TAPE (Supplementary Figure S9) and CSx-S (Supplementary Figure S10). Both MODE and TAPE predicted a greater fraction of granule neuron progenitor cells (GNPs) and rhombic lip (RL) in SHH-MB and G3/4-MB, respectively (Supplementary Figure S11). G3-MB tumors had the most abundant RL fraction compared to the other two subgroups, while G4-MB tumors were predominantly enriched in Unipolar Brush Cells (GlutaCN/UBC), with p-values shown in Supplementary Figure S11 and consistent with the results in previous studies [8, 48]. The higher abundance of GNPs in the SHH subgroup, quantified by MODE and TAPE, further demonstrated the role of GNPs in SHH pathway activation during MB tumorigenesis [51]. In contrast, CSx-S mode did not recover the expected abundances of GNP in SHH-MB and RL in G3-MB. Notably, the cellular fractions resolved by MODE demonstrated a previous finding [8] that GlutaCN/UBC origin cells were significantly more abundant (*p <* 10^*−*4^ in DNAm; *p* = 0.0002 in RNA) than RL in G4-MB tumors (Figure 5b), which was incorrectly quantified by TAPE.

**Fig. 5.**
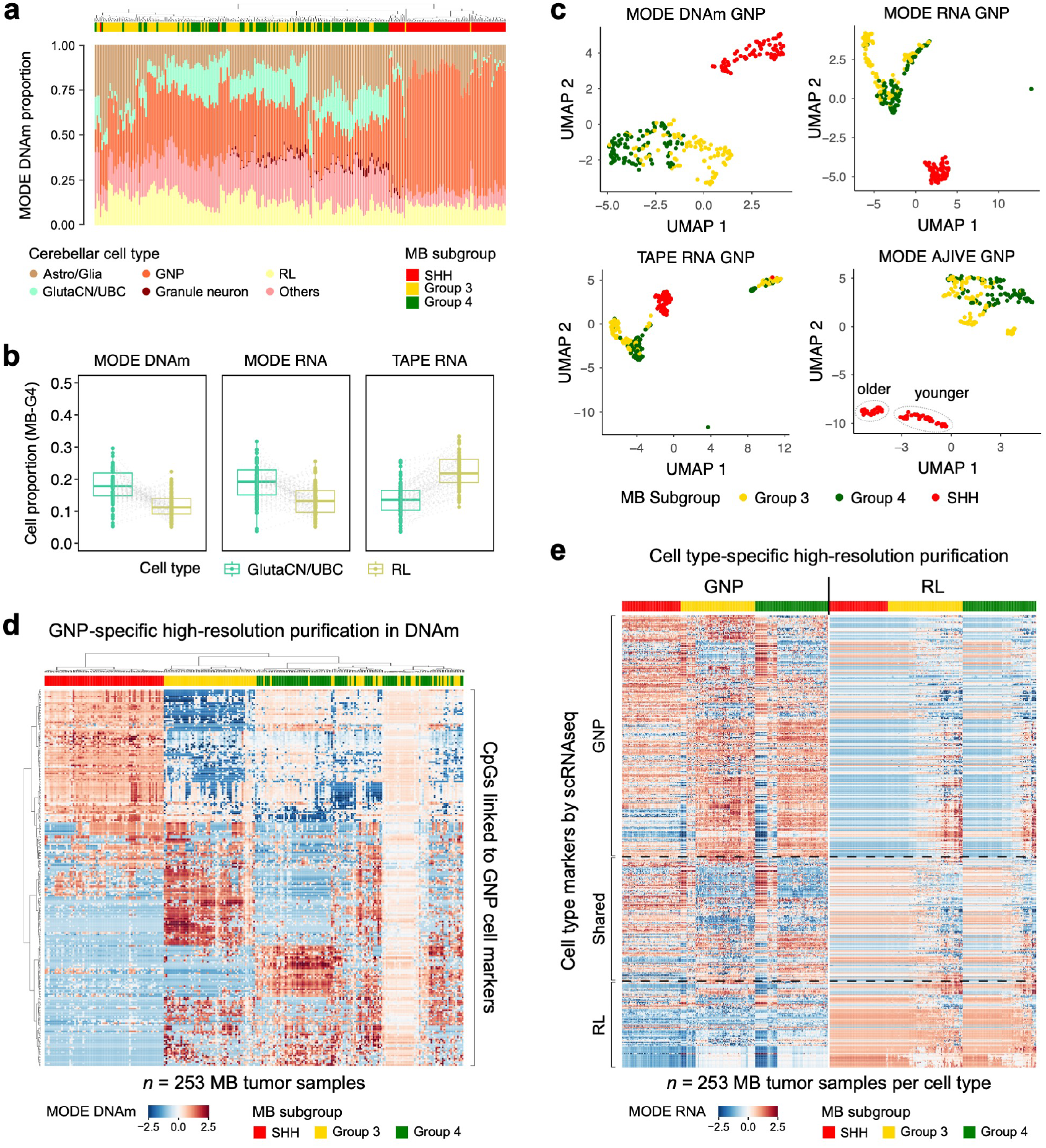
Multi-omic dissociation of pediatric medulloblastoma tumors with human fetal cerebellum scRNA-seq reference. **(a)** Origin cell proportions in bulk MB methylomes quantified by MODE. **(b)** MODE (DNAm, RNA) and TAPE (RNA) proportions of GlutaCN/UBC and RL in G4 MB tumors. **(c)** UMAPs of GNP-specific high-resolution profiles purified by MODE in DNAm, RNA and by TAPE in RNA; UMAP of AJIVE-integrated GNP-specific multiomes purified by MODE. **(d-e)** MODE-predicted MB tumor DNAm (d) and gene expression (e) mapped to GNP or RL lineages for the marker genes identified in scRNA-seq data of human embryonal cerebellum.

The digitally purified MB tumor methylation and gene expression also demonstrated superior power of MODE. DNAm beta values for select CpGs and expression levels for the lineage-specific genes in MB tumors were successfully assigned to the GNP subpopulation and visualized by UMAP (Figure 5c), including a joint visualization of MODE purified multi-omic profiles using the AJIVE algorithm [42, 52]. Surprisingly, the AJIVE-UMAP visualization of MODE multiome (Figure 5c, Supplementary Figure S13) revealed a subcluster of SHH-MB patients who were diagnosed at older age and classified as the delta (i.e., adults) or alpha (i.e., adolescents and children) subtypes [53, 54], driven by the underlying GNP-specific epigenomic heterogeneity. These SHH subtypes were not profoundly visible in the UMAP of bulk data, implying that epigenomic signatures heralding a delayed evolution of SHH-MB tumor may arise from the GNP origin. Moreover, the GNP-specific DNAm profiles predicted by MODE (Figure 5d) showed clear discrimination between SHH and G3/4 with differentially methylated CpGs. This epigenomic signal in the proximity of GNP cell markers aligns with the correlation between neuronal progenitors and G3/4-MB tumors reported in previous transcriptomics studies [55, 56], but rigorous experiments and more evidence are needed to uncover the role of DNA methylome in the differentiation and evolution of neuronal progenitors into MB cancer cells.

In the gene expression profiles purified by MODE, GNP-only markers are highly expressed within the GNP subspace projected to MB tumors, while RL-only markers show overall higher expression in the RL subspace (Figure 5e). On the other hand, CSx predicted the expressions of 273 genes in GNP and 2116 genes in GlutaCN/UBC, but the UMAP clusters of purified samples did not align with MB subgroups (Supplementary Figure S14). We further assessed the DEGs for MB subgroups based on purified data for GNP and/or RL markers, identifying 446 GNP marker genes and 175 RL marker genes differentially expressed (*p <* 0.05) among MB tumor subgroups derived from each developmental cell state (Figure 5e). Several genes previously reported as subgroup-specific drivers or tumor lineage markers, such as *PTCH1, PDE1C, ST18* in SHH [9, 49] and *OTX2, EOMS, LMX1A* in G3 [9, 57], were recovered and detected as DEGs exclusively in GNP and RL, respectively. This result aligns well with the fact that GNP and RL are the embryonal origins for SHH and G3 MB tumors.

### TCGA glioblastoma RNA-seq and DNAm

To further investigate the deconvolution power and robustness in median-size cohorts, we applied MODE and TAPE to the TCGA Glioblastoma multiforme. This dataset contains bulk RNA-seq and DNA methylation paired by the tumors from *n* = 64 GBM patients. We used a preprocessed and annotated scRNA-seq data from GBM patients in an external cohort [6, 34] as reference to resolve the microenvironment of bulk transcriptome-methylome with six cell types (astrocyte, GBM cancer cell, myeloid, neuron, oligodendrocyte, and T cell). There are 2115 DEGs identified as cell markers from this scRNAseq data [34]. We selected the top 10000 variation CpGs and then mapped the probes to DEGs using MethylMix [50], resulting in 965 CpGs. The clinical data of patients were collected from cBioportal [58]. The Kaplan Meier (KM) curve and log-rank test analysis were conducted based on the high- and low-cell abundance groups stratified by the binarized estimated cell proportions.

The scRNA-seq reference data was profiled from two GBM patients with identifiers SF3448 and SF12460, individually, as visualized in Figure 6a. Both MODE and TAPE use the same default sparsity parameter to evaluate the impact of reference data on each method. Obviously, the confidence intervals per cell type in (Figure 6b) implies significant difference between methods and molecules in the context of prediction uncertainty, validated by the Fligner-Killeen test comparing variances (with p-values shown in Supplementary File 4). Specifically, MODE yields smaller variation of cellular fractions in either DNAm or RNA compared to TAPE, especially in astrocyte and GBM fractions (*p <* 10^*−*4^). The neuron cell fraction was stably recovered by MODE from the bulk multi-omics data using either scRNA-seq reference, while TAPE failed to quantify this subpopulation in many target tissues using reference SF34348. On the other hand, the estimated abundance of oligodendrocyte in RNA was not ideal in either method, but MODE was able to recover this cell state in the bulk DNAm profiles. The overall distribution of the resolved tumor microenvironment based on either reference aligns with our hypothesis that MODE was not severely affected by the cell annotation quality in scRNA-seq reference because of the target tissue-recovered reference panel used in the training process.

**Fig. 6.**
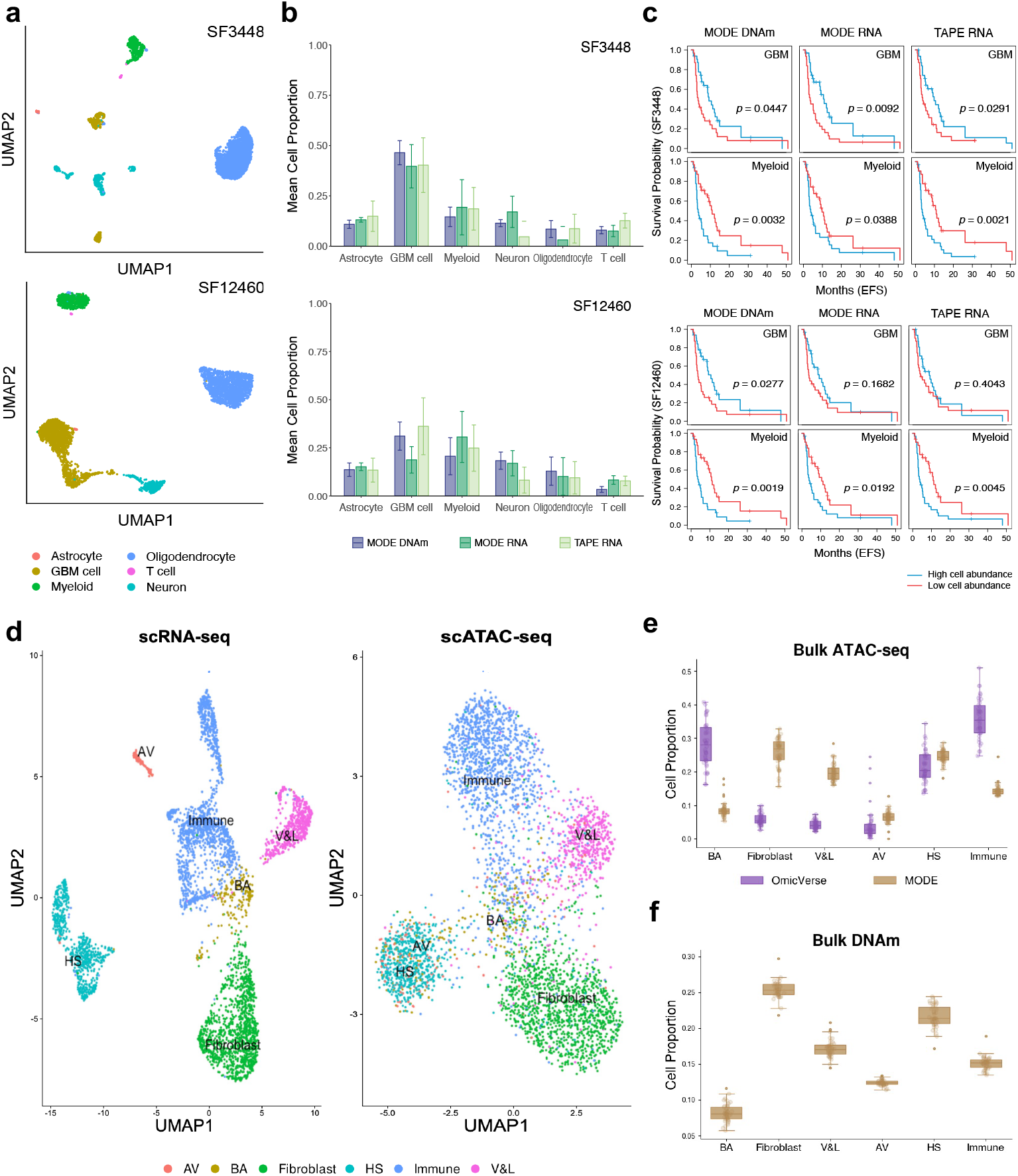
Dissociation of TCGA bulk multi-omics data with different scRNA-seq references. **(a)** Uniform Manifold Approximation and Projection (UMAP) visualizations of an external scRNA-seq reference data collected from two GBM tumors (patients). **(b)** The mean cellular compositions estimated by MODE (DNAm, RNA) and TAPE (RNA) with single cell reference in **(a). (c)** TCGA GBM patients’ survival outcomes stratified by binarized cell proportions. **(d)** A breast cancer tumor transcriptome (left) and epigenome (right) paired by single cells with annotations determined by RNA-seq clusters and refined by ATAC-seq clusters. **(e-f)** TCGA BC epithelial and stromal cells compositions resolved by MODE and OmicVerse.

We further examined the predicted microenvironment composition by linking it to the survival outcomes of patients who were classified as high vs low cell fraction subgroups based on the median abundance per cell type. Both MODE and TAPE uncovered that higher myeloid abundance was associated with poor event free survival (EFS) across different deconvolution signature matrices (see Supplementary Table S1), being consistent with a recent finding about myeloid in GBM [59]. For the reference SF3448, MODE RNA proportion of GBM tumor cells (*p* = 0.0092) presented prognostic detection power comparable or slightly higher than TAPE RNA result (*p* = 0.0291) (Figure 6c). Meanwhile, the GBM proportion in methylome estimated by MODE with reference SF12460 revealed clear EFS stratification among patients (*p* = 0.0277), while TAPE cannot identify this difference in RNA with the same reference (*p* = 0.4043). This result also implied that higher tumor purity in GBM tissues may delay the recurrence or progression of this disease. Similarly, by using the same reference of SF12460, the neuron and T cell fractions in DNAm showed better stratification of EFS (Supplementary Table S1), individually, which was not observed in TAPE RNA fractions or previous GBM deconvolution analyses. The log-rank test results for other cell states and overall survival outcome were shown in Supplementary Figure S15 and Table S1. In summary, by recovering the pure cell bulk methylome from target tumors, MODE unraveled the tumor microenvironment accurately with higher prognostic detection power than TAPE.

### TCGA breast cancer ATAC-seq and DNAm

We used single-cell paired gene expression and chromatin accessibility from human breast cancer (BC) tumor tissue [60] to construct the scATAC-seq reference for deconvoluting n=42 BC tumors in the TCGA cohort profiled by bulk ATAC-seq [61] and DNAm [62] assays. We used the mammary epithelial and stromal cell markers found in a previous BC study [60] to annotate this scMultiome reference. Details of cell labeling and cell marker identification in scATAC-seq reference were described in Methods. The tissue-paired bulk ATAC-seq and DNA methylation data were filtered to retain peaks and CpGs mapped to the 1019 cell marker genes identified from scATAC-seq. The counts in the bulk ATAC-seq were aggregated based on the mapping between peaks with genes, while 720 CpGs were selected from the 10,000 most variable CpGs based on the mapping to cell marker genes, similar to the TCGA GBM DNAm data preprocessing. To train multimodal autoencoder in MODE, we used the JNMF purified output to construct pseudo bulk DNAm and scATAC-seq reference to build pseudo bulk ATAC-seq. In this cohort, we compared our method to OmicVerse because this tool was one of the top performers in real pseudo bulk ATAC-seq deconvolution (Figure 2c) and used a variational autoencoder framework. Both tools were run without fine-tuning.

In the originally annotated scMultiome (Supplementary Figure S16), there were a substantial number of basal/myoepithelial (BA) cells mixed with the clusters of Immune, Fibroblast, and vascular & lymphatic (V&L) cells in scATAC-seq, although these BA cells were well clustered in scRNA-seq. Hence, we reannotated the ambiguous cells based on the scATAC-seq Seurat cluster membership (Figure 6d). The results of MODE and OmicVerse using this reannotated reference were visualized in Figure 6e-f. Notably, the lineage cell and tumor microenvironment compositions resolved by MODE (either ATAC-seq or DNAm) were consistent with the cellular subpopulations in the scRNA-seq and CyTOF single cell proteomics of BC tissues [60], whereas hormone-sensing (HS) epithelial cells and Fibroblast were the most abundant cell states. On the other hand, OmicVerse deconvolution in bulk ATAC-seq (6e) did not quantify the epithelial or stromal cell types at the expected abundances, whereas Fibroblast was *<* 10% and BA was more abundant than HS.

To assess how each method responds to the quality of scATAC-seq reference, we compared ATAC-seq cell proportions predicted by OmicVerse and MODE using both the original and reannotated single-cell references. To reduce the impact of DNAm feature selection on MODE performance, we included all the CpGs (*>* 20, 000) mapped to the 1019 cell marker genes identified in scATAC-seq in this sensitivity analysis. One notable difference is that OmicVerse yields a worse (increased) BA proportion post reannotation, whereas MODE ATAC-seq result showed an improvement (decrease) in BA with median proportion changed from 12.3% to 7.5% (Supplementary Figures S17-S18). This indicates MODE’s reasonable responsiveness to the improved single cell reference quality. The Fibroblast abundance remained low in OmicVerse after reannotation, while MODE Fibroblast in ATAC resolved by the low-quality reference was the most abundant cell type and became less abundant and closer to HS fraction after using the reannotated reference. This is consistent with the results in Figure 6e. Overall, OmicVerse appeared to be more sensitive to the changes in single-cell reference with more visible shifts in multiple cell types, while MODE cell fractions distribution in either modality was stable across different single cell references.

## Discussion

We developed a multimodal deconvolution method MODE that captures cellular processes between molecular modalities, generalizable to any tumor and normal tissue dissociation. Through JNMF initialization, MODE recovered the purified profiles from bulk epigenome or proteome and provided a personalized reference panel to build pseudo bulk samples for training. The multimodal autoencoder network in MODE jointly quantifies the cell abundances in RNA and non-RNA molecules through a shared cell count representation and yields cell-type specific profiles per modality. This is an efficient tool that only requires 5 minutes to dissociate 40 pairs of bulk multi-omics data with approximately 1000 features (e.g., genes, CpGs, proteins) in each modality, using a Lenovo Legion 5i laptop with 32GB RAM.

Our method is distinguished with four key advantages compared to the existing single omics deconvolution methods. (1) MODE can accurately estimate non-RNA cell proportions by multimodal joint learning with single cell reference in only one modality, without the requirement of cross-modality linked features. (2) The multimodal autoencoder network is trained on the simulated pseudo bulk multiome samples, of which the non-RNA pseudo bulk is built on top of the purified data recovered from target samples. This approach not only bridges the gap between single cell reference and target bulk data, but also incorporates realistic between-sample heterogeneity into the training data. (3) MODE introduces the proportional shift strategy in Step 2 to effectively link RNA and non-RNA cellular composition in the same sample to enable integrative deconvolution. (4) MODE can recover rare cell states to decipher the complex tumor microenvironment and potentially identify cts-signatures predictive of disease onset and prognosis.

MODE does not require fine-tuning for the estimation of cellular composition based on the results in Supplementary File 1 and the other simulation datasets. However, these hyperparameters are still critical for the accuracy of purification. Particularly, the JNMF gradient descent optimization step size *s, p* and convergence threshold *ϵ* coupled with the autoencoder epoch and batch size have substantial impact on the purified multi-omic data. Users can rerun our pipeline with various configurations and use a validation cohort of multi-sample single cell data profiled from similar tissues to construct real pseudo bulk samples (similar to Figure 2b) and then merge them with the target data, using the ground truth CTS profile to select the optimal configuration. Moreover, if a subset of tissues are profiled by both single-cell and bulk assays, the tissue-paired single-cell data can be used as the ‘ground truth’ to select the optimal hyperparameters. On the other hand, if none of the target bulk tissues is paired with a single cell profile and a multi-sample single cell validation cohort is unavailable, an alternative solution is to use 80% of the in-silico samples generated in Step 2 to pre-train the autoencoder and then merge the target bulk samples with the remaining 20% in-silico pseudo bulk samples as an augmented target cohort. The parameters minimizing the difference between predicted and ground truth cell type proportions for the 20% training samples is the optimal setting.

The primary goal of MODE is to provide deconvolution and purification for non-RNA or a modality where single cell reference and cell marker signatures are lacking. The simulation results also implied that MODE did not significantly outperform the state-of-art deconvolution methods (e.g., BayesPrism, CIBERSORTx) in RNA-seq data. Further, the current version of MODE does not provide deconvolution of triple-modality omics data (e.g., RNA-seq, ATAC-seq, proteomics paired by tissues). But this type of analysis is feasible if users revise the algorithms by adding another modality in JNMF and autoencoder, individually. It should be noted that the performance of a revised algorithm on triple-omics requires additional benchmarking experiments because of the increased dimensions in JNMF and autoencoder parameters. We will address this challenge in future work by developing a tool generalizable to triple-modality data for bulk and spatial omics.

MODE improved both cell abundance quantification and high-resolution digital purification of bulk multiome compared to multiple baseline methods, being a framework generalizable to various omics data. In the current work, we assume the sequencing depth or batch effects in reference data to be moderate compared to the profound biological heterogeneity between single cell reference and target bulk samples. If there are strong technical artifacts within the reference data, we can either use a single-batch reference to avoid striking batch effect or use a tool that aims to normalize reference transcriptome size in deconvolution (e.g., ReDeconv [63]) before running MODE. One possible extension to MODE is to enhance the consistency in digital purification by leveraging and imputing an external scMultiome reference. Our current algorithm (Step 2) constructs the pseudo bulk multi-omics data from an external scRNA-seq data and the tissue-recovered non-RNA signature matrices, without imputing the missing features in scRNA-seq. Hence, recovering the undetected features in single cell reference may increase the power of predicting individualized ctsMultiome in future. Another potential innovation built on MODE is to resolve the spatial and cellular landscapes in tissues by integrating bulk multi-omic profiles, whole slide imaging, and a spatial transcriptomics reference. A recent research reconstructed the spatial landscape of malignant cells in breast cancer bulk transcriptome via graph attention neural network [64]. We may incorporate this approach in the MODE framework to predict multimodal cellular landscape per spot, as well as identify the regional biomarkers not detectable in current spatial technologies. Together, we will be able to elucidate the underlying mechanisms for disease with cross-modality segmentation of biospecimens.

## Methods

### The MODE framework

For each tumor tissue, we measure the mixed-cell bulk expression of gene *g* (*g* = 1, …, *m*_1_) and the abundance of feature *h* (*h* = 1, …, *m*_2_) in the other omics (e.g., CpG sites, peaks for chromatin accessibility, proteins), respectively, denoted by *y*_1*g*_, *y*_2*h*_. The latent individualized measurements for cell type *c* (*c* = 1, …, *k*) are denoted by *x*_1*gc*_, *x*_2*hc*_. The modality-specific cellular fractions are *θ*_1*c*_, *θ*_2*c*_, determined by the common tissue-specific cell counts (fractions) *p*_*c*_ and source-specific cell size factors *s*_1*c*_, *s*_2*c*_. That is 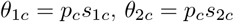. Thus, the paired bulk multi-omic data are described as

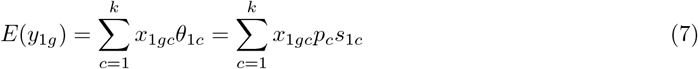

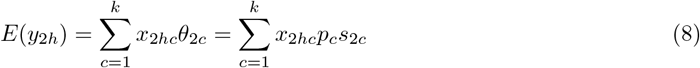

We denote 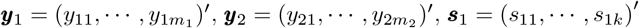 and ***s***_2_ = (*s*_21_, *· · ·, s*_2*k*_)′.

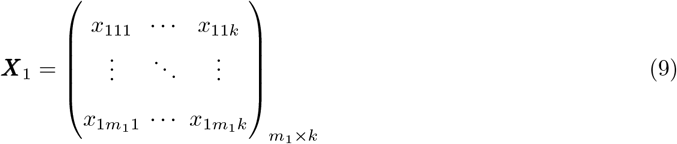

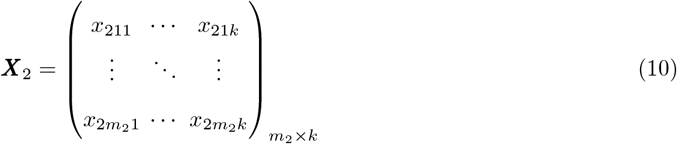

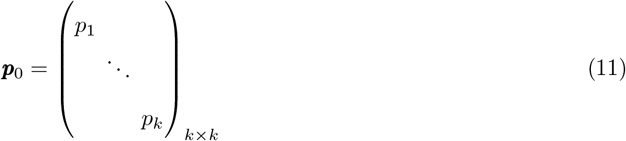

#### Step 1: Initialize ctsMultiome from target tissue

We deploy the Joint Non-negative Matrix Factorization model to pre-purify non-RNA profiles, which serves as reference data to generate large-scale pseudo bulk samples used in the training step. We optimize high-dimensional parameters ***η*** = *{****X***_1_, ***X***_2_, ***p***_0_, ***s***_1_, ***s***_2_*}* by minimizing the *L*_2_ loss function:

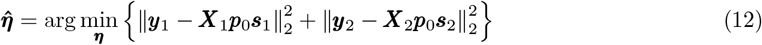

subject to min*{****η****} ≥* 0.

To eliminate the impact of diverse multi-omics data scales on JNMF optimization, we used logarithmic to transform RNA-seq, ATAC-seq, and MS proteomics data, while the DNAm beta values were transformed by negative logarithmic. This step integrates the observed bulk multiome with shared cell counts to pre-estimate the cell-type-specific profiles in each molecular source, with details in our previous work [42]. The parameters in JNMF are randomly initialized based on the distribution of input bulk multiome and the hypothesized RNA cell counts extracted from previous experiments. This is a reference-free approach that initializes the personalized CTS non-RNA profiles without using modality-matched single cell reference.

#### Step 2: Prepare pseudo-bulk multiome for training

Pseudo-bulk data that imitates the target bulk profile can be generated by aggregating single cell expression data with a given cell type proportion. We proposed omics-joint pseudo-bulk simulation to connect the different modalities of matched samples by introducing a proportional-shift strategy. The original RNA-seq cell-type proportion could be generated using Dirichlet distribution. Then, a proportion-shift is applied to the original RNA proportion by multiplication with a uniform distribution. Proportional-shift can create the original proportion of non-RNA-seq omics while preserving similarities between the proportions of different molecular sources.

##### RNA-seq pseudo bulk samples

A preprocessed scRNA-seq gene count matrix with cell type labels was used to generate the artificial training samples of RNA-seq. The original RNA-seq cell type proportions were first generated using the Dirichlet function from the Python package NumPy without *a priori* parameters, i.e., the fraction of each cell type is assumed at 1*/k*, similar to the counterpart in TAPE [28]. Thus, the mean cellular composition across all the training samples are balanced for the cell types of interest. To address the sparsity of training samples heavily biased toward a certain cell type or missing specific cell types, some of the simulated RNA-seq cell type proportions were randomly knocked out and set at zero to build a sparse composition matrix that served as updated ground truth. Next, we determine the exact number of cells to sample from each cell type based on the total number of cells available in the scRNA-seq reference and the ground truth cell type fractions per sample. We then randomly select the appropriate number of cells for each cell type. Finally, the RNA-seq pseudo-bulk sample is produced by aggregating gene expression values across selected single cells.

##### Non-RNA pseudo bulk samples

The generation of training data in the other omics leverages the pre-purified profiles output from JNMF in Step 1. The non-RNA cellular fractions were shifted from the above balanced RNA-seq cell type proportions by multiplying a random shifting factor sampled from the Uniform distribution *U* (0.9, 1.1) using the NumPy package, randomly scaling the proportions between 90% and 110% of the original value. Sparsity was introduced to this shifted non-RNA proportions through the same knock-out procedure used in RNA-seq pseudo bulk generation. Next, we calculate the mean mixed-cell expression (or abundance) for the input features by the weighted sum of pre-purified non-RNA profiles (***X***_2_) per cell type in Step 1: 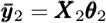, whereas the weights ***θ***_2_ are the cellular composition after random knock-out. To amplify the cross-sample heterogeneity and boost training sample size, a Poisson random generator was used to generate large-scale bulk omics profiles: 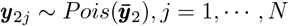. In this data generating process, the pre-purified protein abundance or chromatin accessibility is transformed back to the original (non-log) scale, while the purified DNAm beta values is still at negative logarithmic scale. The number of pseudo bulk samples (*N ≥* 5000) is expected to be larger than the sample size of real target tissues (*n*).

#### Step 3: Multimodal tissue-adaptive dissociation

We propose a multimodal autoencoder to learn the shared molecular signals described by different omics profiles for tissue-adaptive deconvolution. This multimodal autoencoder contains one encoder (*E*_0_) for the concatenated bulk input from RNA and non-RNA profiles and two parallel decoders (*D*_1_, *D*_2_) for the reconstruction of different bulk omics data. The concatenated bulk expression profiles are reduced to latent representation ***p***_*j*_ ∈ ℝ^*k*^ (*j* = 1, *· · ·, N*), representing the shared cell counts fractions per training sample, through a five-layer fully connected neural network. We use the encoder to predict cell counts fractions for all the training samples 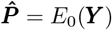, whereas ***Y*** = (***Y*** _1_, ***Y*** _2_), ***Y*** _1_ is a *N ×m*_1_ matrix for the pseudo bulk RNA-seq gene expression profiles and ***Y*** _2_ is a *N × m*_2_ matrix for the simulated non-RNA bulk profiles at training stage. The bulk multi-omic input data are sample-matched with *m*_1_ and *m*_2_ features, respectively. The CELU activation function is used for the first four hidden layers in the encoder, formulated as CELU(*x*)=max(0,*x*)+min(0,*αe*^*x/α−*1^).

We devise the following two-stage procedure to optimize the encoder and decoder parameters per target tissue. In the training stage, we use pseudo-bulk samples of both omics to train the entire MODE framework by minimizing the deconvolution loss *L*_*d*_ and the reconstruction loss *L*_*r*_. The loss functions are defined as:

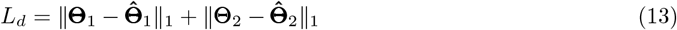

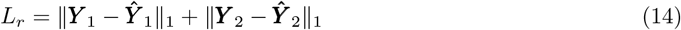

where **Θ**_*i*_, ***Y*** _*i*_ are the ground truth cell proportions and the bulk multi-omic input for *N* training samples. The deconvolution loss calculated between the predicted cell proportions and the ground truth is used to optimize the parameters of encoder and neural network connection layers, while the reconstruction loss between the predicted and the input bulk data is used to simultaneously optimize the connection layers for both decoders and encoder. We use *L*_1_ loss in the calculation of deconvolution loss and reconstruction loss.

In the adaptive stage, we further optimize the encoder and decoders with tissue-specific loss functions and (16). Let 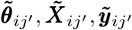 denote the tissue-adapted cell proportions, high-resolution reference panel, reconstructed bulk profiles for modality *i* in target tissue *j*′ (*j*′ = 1, *· · ·, n*) based on the parameters learned in the training stage, ensuring the parameters not over fitted compared to the original estimates derived on pseudo bulk training data. The adaptive stage allows personalized deconvolution with parameters re-optimized to each pair of target bulk samples and further improve the prediction beyond training by minimizing the following loss functions.

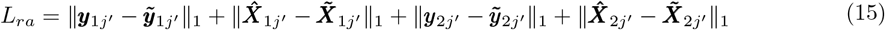

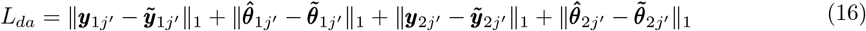

### Simulation data

#### Pseudo bulk RNA-seq and ATAC-seq paired by developing human brain

To design this experiment, we obtained the cell-matched raw count matrix and cell type annotations for 45549 single cells that were simultaneously profiled from 12 human neocortex samples using multi-omic snRNA-seq and snATAC-seq (GSE204684). To effectively leverage the information from both modalities, we adopt the following steps to clean and merge the cells: 1) remove various stromal cell types with smaller population sizes, including endothelial cells, pericytes, and vascular smooth muscle cells (VSMCs); 2) exclude radial glia (RG) and intermediate progenitor cells (IPCs) from the analysis due to the insufficient sensitivity of RNA-seq data in accurately identifying these progenitor populations, as indicated by nearly 20% lower detection compared to ATAC-seq data; 3) merge the medial ganglionic eminence (MGE)-derived and caudal ganglionic eminence (CGE)-derived inhibitory neuron subtypes, as they were not distinguishable in the ATAC-seq clustering. As a result, a total of 42027 cells across nine major neocortical cell types in both modalities were retained for downstream analysis (Figure 2a). Given the significant differences in cell population composition between fetal and non-fetal samples, we designated single cells from one early mid gestation fetal sample, one late mid gestation fetal sample, and one infant sample as reference signatures. Single cells from the remaining seven non-fetal samples were utilized to construct multi-omics pseudo bulk samples. For RNA-seq, each pseudo bulk sample was created by aggregating all single cells in the respective sample. The RNA cell proportions in each sample were calculated by the ratios of cell counts per cell state vs. the total of cells. We shifted the true RNA proportions by random factors generated by the Uniform distribution *U* (0.6, 1.4) to derive the sample-matched true ATAC-seq proportions, achieving a Pearson correlation *r* = 0.97 between two modalities in the ground truth cellular fractions. Next, we sampled 1000 single cells from each non-fetal sample without replacement and aggregated them to construct a pseudo bulk ATAC-seq sample. These seven paired non-fetal pseudo bulk multi-omics samples served as target bulk data for the deconvolution analysis. To identify markers (genes or peaks) for each annotated cell type, we applied the FindMarkers function in Seurat to scRNA-seq and scATAC-seq references, respectively, selecting genes or peaks with Bonferroni adjusted p-value *<* 0.01 and LFC *>* 0.5. We evaluated the deconvolution performance of MODE and competing methods using the same cell type marker list per modality.

#### Simulation design for scMultiome in glioblastoma tumors

We designed a rigorous simulation study to mimic the intra- and inter-tumor heterogeneities in bulk multi-omics data. Herein, we used the real single cell multi-omics data from adult GBM tumor tissues profiled by the scRNA-seq and scATAC-seq platforms, individually, to extract the simulation parameters, downloaded from GEO (GSE17455) and https://db.cngb.org/cdcp/dataset/SCDS0000041. The scRNA-seq data were preprocessed and annotated in a previous analysis [34], while the tissue-matched scATAC-seq data still lacked cell labels. Thus, we identified the anchors between RNA and chromatin datasets from the same biological system by using FindTransferAnchor in Seurat 3.0 [17] and then transferred the cell labels of scRNA-seq to scATAC-seq data via function TransferData. The estimate of gene activity matrix for scATAC-seq was quantified by the total counts within 2kb upstream region of the gene body. The annotated real scATAC-seq data along with the tissue-matched scRNA-seq data were visualized in Supplementary Figure S4a.

Given the ambiguous cell annotation and rare cell types in this single cell data (Supplementary Figure S4a), we first removed the low-quality cells. To introduce randomness in each in-silico tissue and reduce computation time of scDesign3, we randomly sampled 500 cells from five major cellular populations within this real single cell sample to estimate the parameters for cell-specific mean expressions and cell-to-cell heterogeneity (Supplementary Figure S4b), which were later used to generate synthetic cells. The function fit marginal in scDesign3 was employed to extract the feature-wise mean and overdispersion, in which the statistical models were zero-inflated Poisson for scRNA-seq counts and Gaussian for scATAC-seq log-scaled counts. The co-expression network was fitted by Gaussian distribution via fit copula function for each modality. Further, we used the UMAP spatial coordinates extracted by R package uwot from the 500 real single cells as the tensor regression covariates along with the above feature-wise parameters in the simulator function (simu new) to simultaneously generate multiome (gene expression and chromatin accessibility) at cell resolution. This simulator was rerun 200 times to generate a large pool of n=100,000 in-silico single cells of multiome with 542 genes and 888 peaks. To generate each pseudo bulk multi-omics sample, 500 synthetic cells of each run were slightly subset based on the sample-wise cellular composition pre-simulated by Dirichlet distribution and then aggregated as the pseudo bulk admixture gene expression or chromatin accessibility (Supplementary Figure S4c). This procedure ensures disparities between in-silico tissues since the synthetic cells used in different samples are not overlapping within each modality.

The difference across two molecular sources in this experiment is determined by the cellular compositions between paired omics samples. In the generation of pseudo bulk ATAC and RNA cellular compositions, we used the Uniform distribution to control the ‘shift’ in cell proportions from RNA to ATAC. We first simulated the ground truth RNA cell proportions by Dirichlet distribution, with parameters estimated from the reference multi-subject cell counts. In the scenario of small difference, we randomly generated a ‘shifting’ factor (of dimension *K*) for each sample via Uniform distribution *U* (0.9, 1.1) and then multiplied this factor with the RNA proportions followed by sum-to-one normalization, which is the ATAC proportions. For the scenario of large difference, we use *U* (0.6, 1.4) to generate the sample-wise shifting factor, allowing larger difference in ground truth cell proportions across modalities.

#### Evaluation metrics for purified bulk proteome and transcriptome

In each cell type, the biological (or theoretical) difference in the log-scaled gene expressions between two disease groups is defined by the log fold change (LFC) value (e.g., LFC=0, 0.5, 1, 1.5). Specifically, we generate the population mean gene expressions per disease group using Multivariate Normal (MVN) distribution MVN(*µ*, Σ) with group-wise mean parameters *µ*_1_, *µ*_2_, whereas log (*µ*_1_) *−* log (*µ*_2_)=LFC. We set 100 genes in astrocyte and neuron, individually, with LFC ≠ 0 as the ground truth ctsDEG, while the other genes were set at LFC=0 (as ‘null’ genes) per cell type. The mean gene expression parameter is converted back to the original scale by 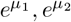 in the remaining steps of generating RNA-seq raw counts, described in our previous work [42, 65]. We evaluated the quality of high-resolution ctsMultiome profiles purified by MODE and TAPE based on the accuracy of downstream ctsDE detection. In the ground truth of astrocyte- and neuron-specific multi-omics expression data, there are 100 features per modality generated as ctsDE for each cell type. For the purified data in each molecular source, features with zero variance across samples were considered invalid and excluded in the downstream DE analysis. We used Wilcoxon rank-sum test to identify DE features in cts-transcriptome and -proteome, individually, with false discovery rate (FDR) corrected by Benjamini-Hochberg (BH) procedure. Features with FDR-adjusted *p <* 0.05 were called as DE. We used sensitivity, specificity, and the F1-score between ground truth vs. predicted labels as metrics to evaluate the power and robustness of DE detection.

#### Preprocessing single cell and bulk multiome in breast cancer

The single cell epigenome and transcriptome (GEO accession: GSM6543828) were paired by cells and split into scRNA-seq and scATAC-seq to create separate Seurat objects, both normalized and clustered using Seurat functions. Marker genes were then identified from the scRNA-seq clusters using Seurat’s FindMarkers() function and matched to a list of major breast cancer cell type markers [60] to assign cell type labels. These annotations were transferred to the scATAC-seq data, which was then grouped by cell type to identify cell type specific marker peaks using FindMarkers(). Peaks were mapped to genes by selecting those with start positions within the gene body or up to 2 kb upstream of the transcription start site. These peaks were then aggregated to gene level by the total accessibility mapped to gene within each cell, resulting in a scATAC-seq reference (1019 genes x 4641 cells) with genes mappable to the peaks in TCGA bulk ATAC-seq data. The peak-gene and CpG-gene mapping in the bulk multi-omics data were downloaded from https://xenabrowser.net/datapages/.

## Competing methods

### CIBERSORTx

CIBERSORTx is a computational framework designed to accurately estimate cell type abundance and cell-type-specific gene expression profiles from bulk RNA-seq data. Building upon its predecessor, CIBER-SORT, it incorporates new functionalities such as cross-platform data normalization and in silico cell purification, enabling the digital isolation of individual cell-type transcriptomes without the need for physical separation.

### Scaden

Scaden is a deep learning-based deconvolution model designed to estimate cell type proportions from bulk RNA-seq data. Its basic architecture is a deep neural network that takes gene count data from RNA-seq as input and outputs predicted cell fractions. The model employs an ensemble approach, combining the predictions of the three best-performing models to produce the final cell type composition estimates.

### TAPE

TAPE is an autoencoder framework designed to accurately predict cellular fractions and cell-type-specific gene expression from bulk RNA-seq data. TAPE demonstrates competitive performance compared to state-of-the-art methods while offering one of the fastest processing speeds on benchmark datasets. TAPE’s tissue-adaptive prediction of cell-type-specific gene expression enables the dissection of bulk gene expression into different cell types.

### scpDeconv

scpDeconv is a deep learning-based method to estimate cell-type proportions from tissue proteomic data, utilizing single-cell proteomic data from similar cohorts as a reference. This approach integrates a mix-up strategy, autoencoder, and domain-adversarial neural network to overcome several challenges in proteomic deconvolution. scpDeconv eliminates the need for extensive simulation datasets by imputing the missingness of low-abundance proteins through its autoencoder component.

### DISSECT

DISSECT is a variational autoencoder algorithm that infers cellular composition and cell-type-specific gene expression profiles from bulk RNA-seq data. During the training phase, DISSECT uses a combination of reconstruction loss and Kullback-Leibler divergence to optimize the model parameters, iteratively updating cell type proportions and cell-type-specific expression profiles to improve deconvolution accuracy.

### BayesPrism

BayesPrism is a Bayesian deconvolution algorithm consisting of four steps. That is inferring joint posterior distributions, estimating cell state-specific parameters, aggregating results to the cell type level, and optionally updating the reference matrix to improve estimates across bulk samples.

### OmicVerse

OmicVerse is a computational tool designed for deconvoluting bulk RNA-seq data using single-cell references. It supports multiple deconvolution algorithms and incorporates quality control, normalization, and visualization features to ensure accurate and interpretable results.

### scSemiProfiler

scSemiProfiler is a deconvolution tool that leverages semi-supervised learning to deconvolute bulk RNA-seq using single-cell RNA-seq reference (partially) paired to the target bulk data. This method digitally dissociates bulk tissues at single cell level through variational autoencoder and adaptive learning, being effective in complex tissues. It addresses challenges such as batch effects and incomplete reference annotations by integrating known and unknown cell types.

## Declarations

### Data availability

- Human cerebral cortex: single-nucleus gene expression and chromatin accessibility data is in GEO under accession number GSE204684.
- GBM tumor tissue-matched single-cell multi-omics: scATAC-seq data is in GEO under accession number GSE174554; scRNA-seq data with cell labels is at https://db.cngb.org/cdcp/dataset/SCDS0000041.
- Human fetal cerebellum organoids: the 10X Genomics scRNA-seq data is available in GEO under accession number GSE150153.
- TCGA adult GBM data: the bulk multi-omics data are available at UCSC Xena and https://gdac.broadinstitute.org/.
- Human breast cancer scMultiome is in GEO under accession number GSM6543828.
- TCGA adult breast cancer data: the bulk multi-omics data are available at UCSC Xena https://xenabrowser.net/datapages/.

### Code availability

- MODE (https://github.com/jsuncompubio/MODE)
- scpDeconv (https://github.com/TencentAILabHealthcare/scpDeconv)
- TAPE and Scaden (https://github.com/poseidonchan/TAPE)
- DISSECT (https://github.com/imsb-uke/DISSECT)
- BayesPrism (https://github.com/Danko-Lab/BayesPrism)
- CIBERSORTx (http://cibersortx.stanford.edu)
- OmicVerse (https://github.com/Starlitnightly/omicverse)
- scSemiProfiler (https://github.com/mcgilldinglab/scSemiProfiler)

## Author contribution

This study was conceived by J.S. and Q.L., who jointly developed the algorithm. Q.L. designed the simulation experiments and real data analyses, interpreted the analyses results, wrote the manuscript, and supervised this research. J.S. implemented the algorithm as pipeline, led analyses in simulation experiments, medulloblastoma, and TCGA GBM, visualized results, contributed to manuscript writing. A.M. ran baseline tools in simulation study, independently processed human BC scMultiome and analyzed TCGA BC data, contributed to manuscript writing. W.Z. contributed to deep learning methodology discussion, simulation experiment design, and manuscript editing. T.L., A.B. and Y.P. contributed to simulation study programming, downstream statistical analyses and visualization. K.S. and P.N. provided the data in pediatric medulloblastom and interpreted the analysis results. A.O., G.R., P.N. supervised clinical and/or molecular data collection in pediatric medulloblastoma. All authors reviewed the manuscript.

## Additional Files

**Supplementary File 1 — Results of simulation on pseudo bulk RNA-seq and ATAC-seq**

**Supplementary File 2 — Supplementary Figures and Tables**

**Supplementary File 3 — Simulation results of bulk RNA-seq and mass spectrometry proteomics**

**Supplementary File 4 — Fligner-Killen test comparing variation of deconvoluted TME in TCGA GBM**

