## Supplementary material for "MODE: high-resolution digital dissociation with deep multimodal autoencoder": Supplementary File 2.pdf

Jiao Sun[#], Ayesha A. Malik[#], Tong Lin, Ayla Bratton, Yue Pan, Kyle Smith, Arzu Onar-Thomas, Giles W. Robinson, Wei Zhang, Paul A. Northcott, and Qian Li[*]

#### Contents

**1 Developmental human brain RNA-seq and ATAC-seq** **2**
1.1 Concordance of JNMF purified ATAC with scATAC-seq ground truth . . . . . . . . . . . 2
1.2 Results of scSemiProfiler . . . . . . . . . . . . . . . . . . . . . . . . . . . . . . . . . . . . . . . 2

**2 In-silico gene expression and chromatin accessibility in glioblastoma tumors** **4**

**3 In-silico bulk transcriptomics and proteomics** **6**

**4 Application to pediatric medulloblastoma** **8**

**5 TCGA glioblastoma** **13**

**6 TCGA breast cancer** **14**

### 1 Developmental human brain RNA-seq and ATAC-seq

#### 1.1 Concordance of JNMF purified ATAC with scATAC-seq ground truth

To illustrate that JNMF (MODE Step 1) does not fully recover cellular information from target tissues and does not lead to overfitting in MODE Steps 2-3, we compared the output from JNMF with the ground truth cell proportions and cell-type-specific (CTS) accessibility profiles. The Pearson correlation coefficient (PCC) for JNMF prediction vs. ground truth cell proportions was $< 0.6$ for both ATAC-seq and RNA-seq, being worse than the final predition by MODE autoencoder. Further, the ground truth CTS profile for each target pseudo bulk sample is the sum of peak counts per cell type. The input admixture ATAC data for JNMF was at log scale. Hence, the purified data was transformed back to the original scale (by exponential ) to perform concordance evaluation. A large proportion of the purified values were highly correlated with ground truth, although there were inflated errors (or biases) in each sample. PCC for each sample are shown in each scatter plot below, while the PCC per cell type per sample is reported in Supplementary File 1 under spreadsheet 'JNMF_purified_ATAC_PCC'. The inflated scale of outliers in JNMF purified data does not affect the pre-training of MODE autoencoder, because the pseudo bulk training samples were simulated based on the mean parameter of Poisson and then rescaled by min-max to the range of (0,1) before running autoencoder.

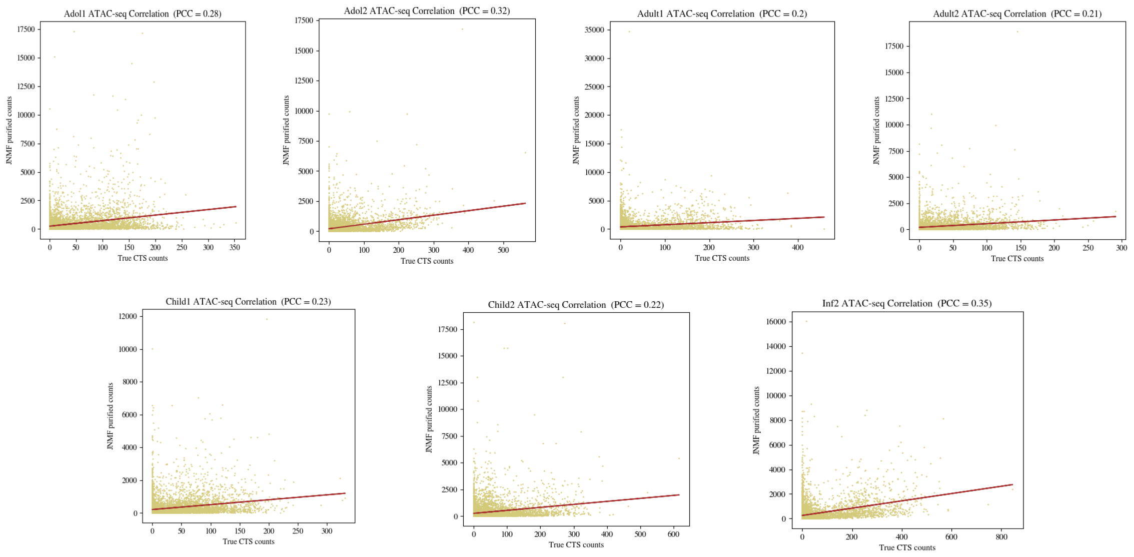


Figure S1: **Pearson correlation for JNMF purified ATAC and ground truth CTS ATAC.**

#### 1.2 Results of scSemiProfiler

The single cell reference samples and pseudo bulk target samples for running scSemiProfiler are described below. For the single cell data of n=2 fetal and n=1 infant neocortex samples, we randomly divided the cells per cell state into 20 groups and then constructed n=20 pseudo bulk samples by aggregating the cells per group. Next, the original n=7 pseudo bulk samples built from postnatal neocortex tissues were merged with the (in-silico) samples from Step 1 as a target cohort for deconvolution, whereas 20 out of 27 samples were overlapped between single cell reference and target bulk data. The predicted cell proportions of the $n = 7$ pseudo bulk samples not overlapping in single cell reference samples are visualized in scatter plots, with evaluation metrics shown in each plot.

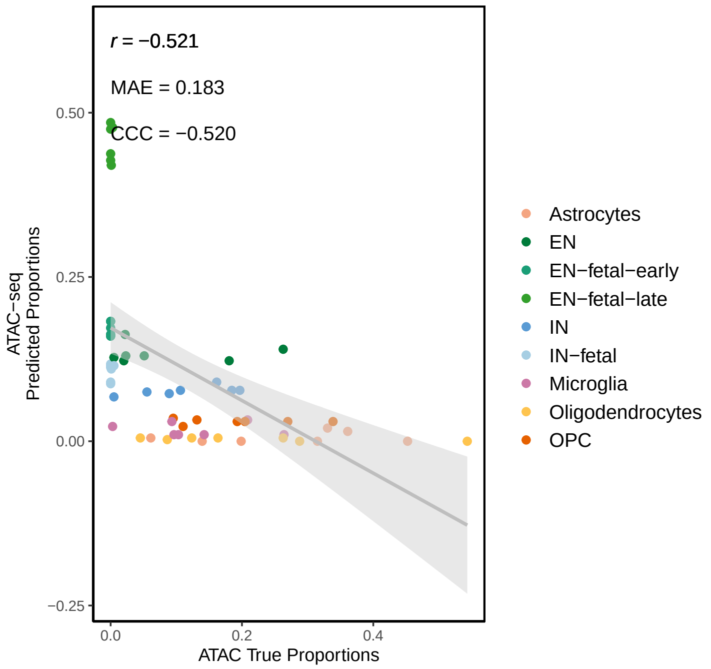


Figure S2: **Scatter plot for scSemiProfiler deconvolution in pseudo bulk ATAC-seq.**

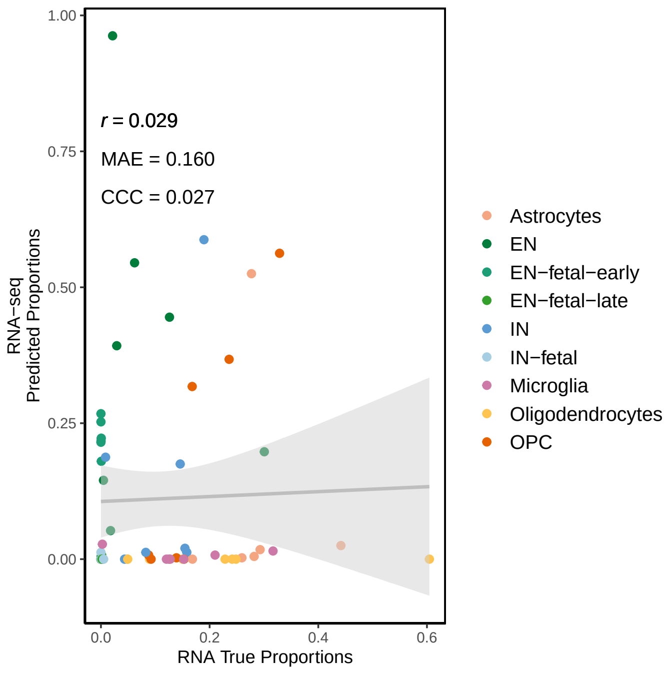


Figure S3: **Scatter plot for scSemiProfiler deconvolution in pseudo bulk RNA-seq.**

#### 2 In-silico gene expression and chromatin accessibility in glioblastoma tumors

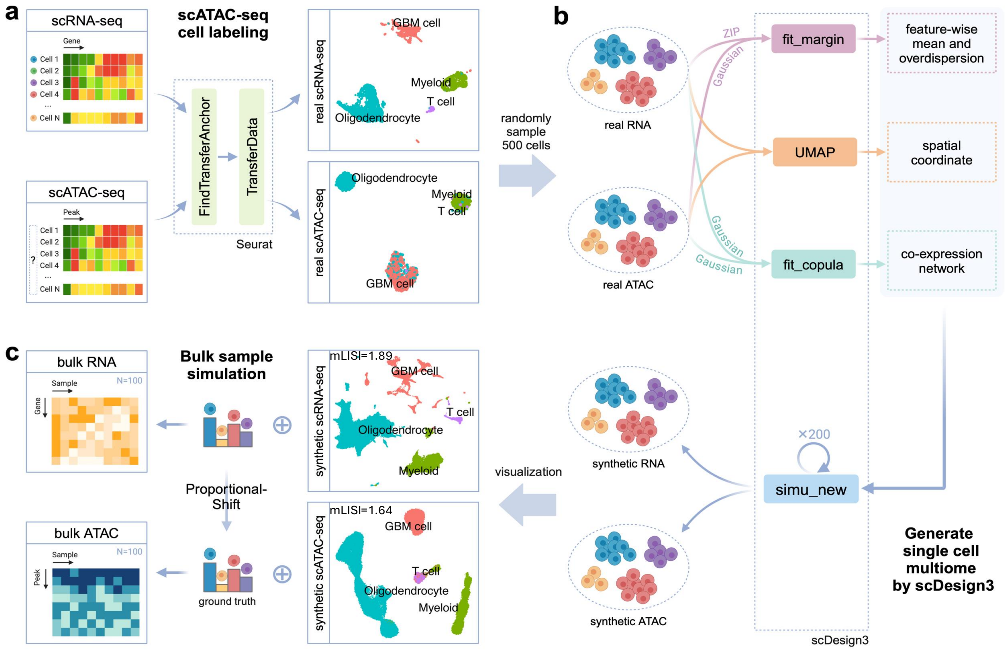


Figure S4: **Simulation of RNA-seq and ATAC-seq paired by tumor tissues.** **(a)** Transfer scRNA-seq cell labels to scATAC-seq by Seurat. **(b)** Synthetic single cell multiome generation by scDesign3. **(c)** Simulated pseudo bulk gene expression and chromatin accessibility, with mean local inverse Simpson's Index (mLISI) shown in the UMAP.

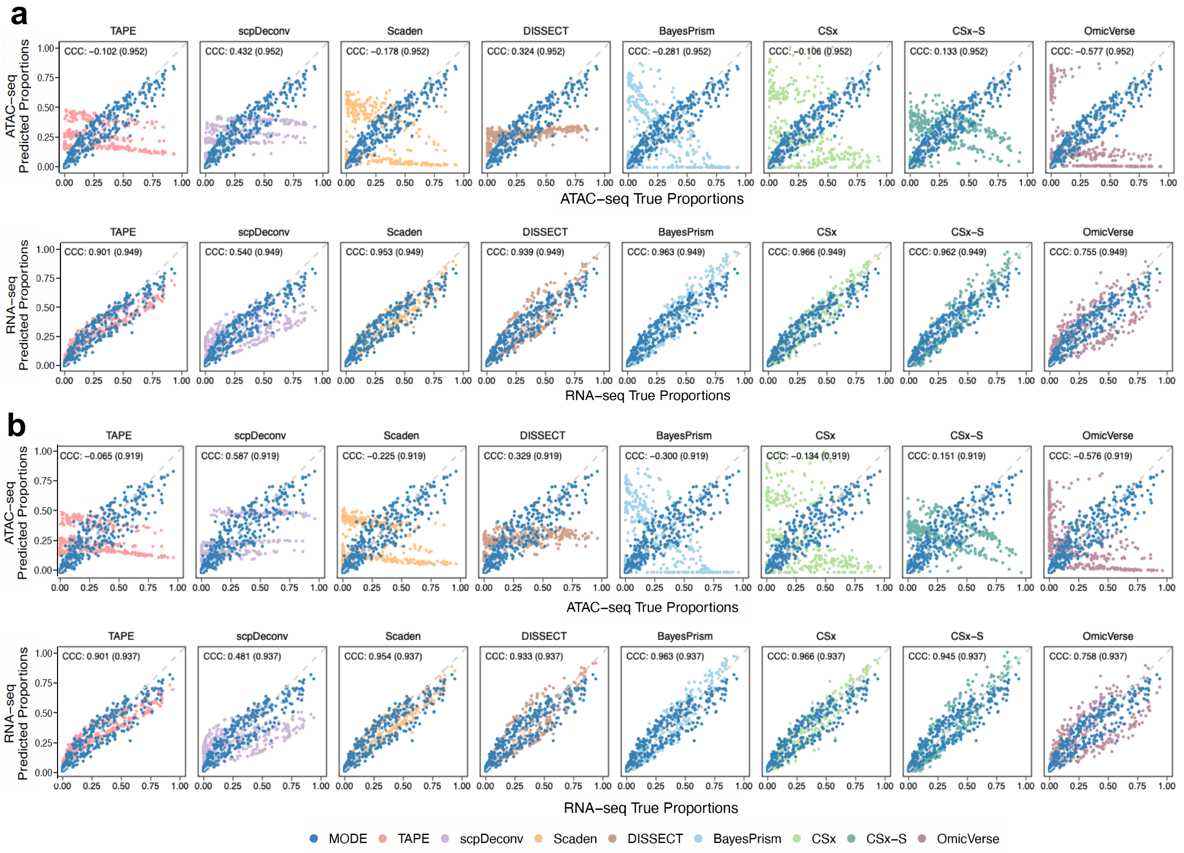


Figure S5: **Performance of competing methods on in-silico RNA-seq and ATAC-seq paired by GBM tumors. (a, b)** Scatter plots of true (x axis) and predicted cell-type proportions (y axis) by deconvolving tumor simulated bulk gene expression and chromatin accessibility with either scRNA-seq or scATAC-seq from GBM tumor as reference. There are two scenarios representing small (**a**) and large (**b**) difference between molecular sources. The CCC between predicted proportion by each competing method and the ground truth is shown at the top of each plot, with MODE result in parentheses.

### 3 In-silico bulk transcriptomics and proteomics

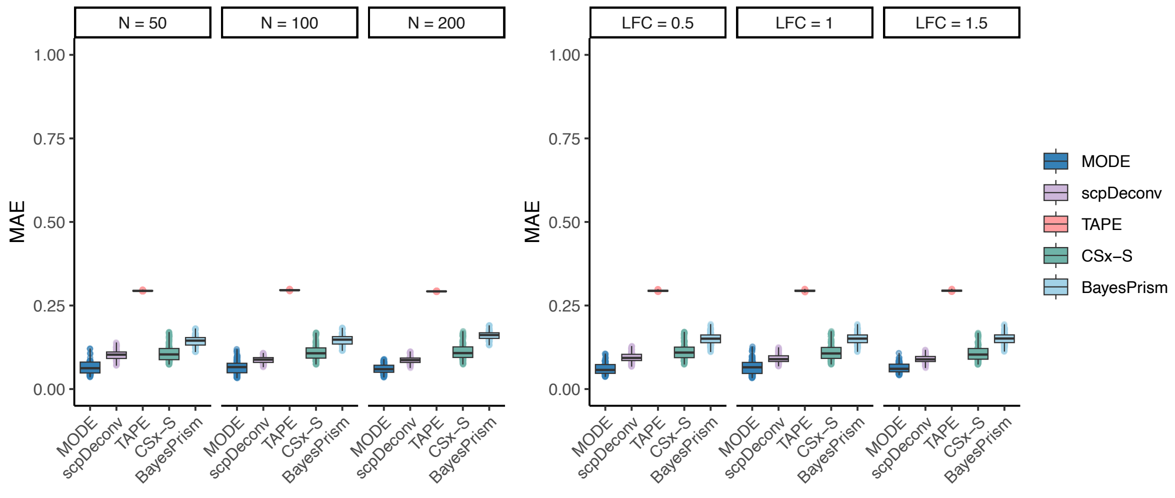


Figure S6: MAE of MODE, scpDeconv, TAPE, CSx-S, and BayesPrism on bulk proteomics deconvolution.

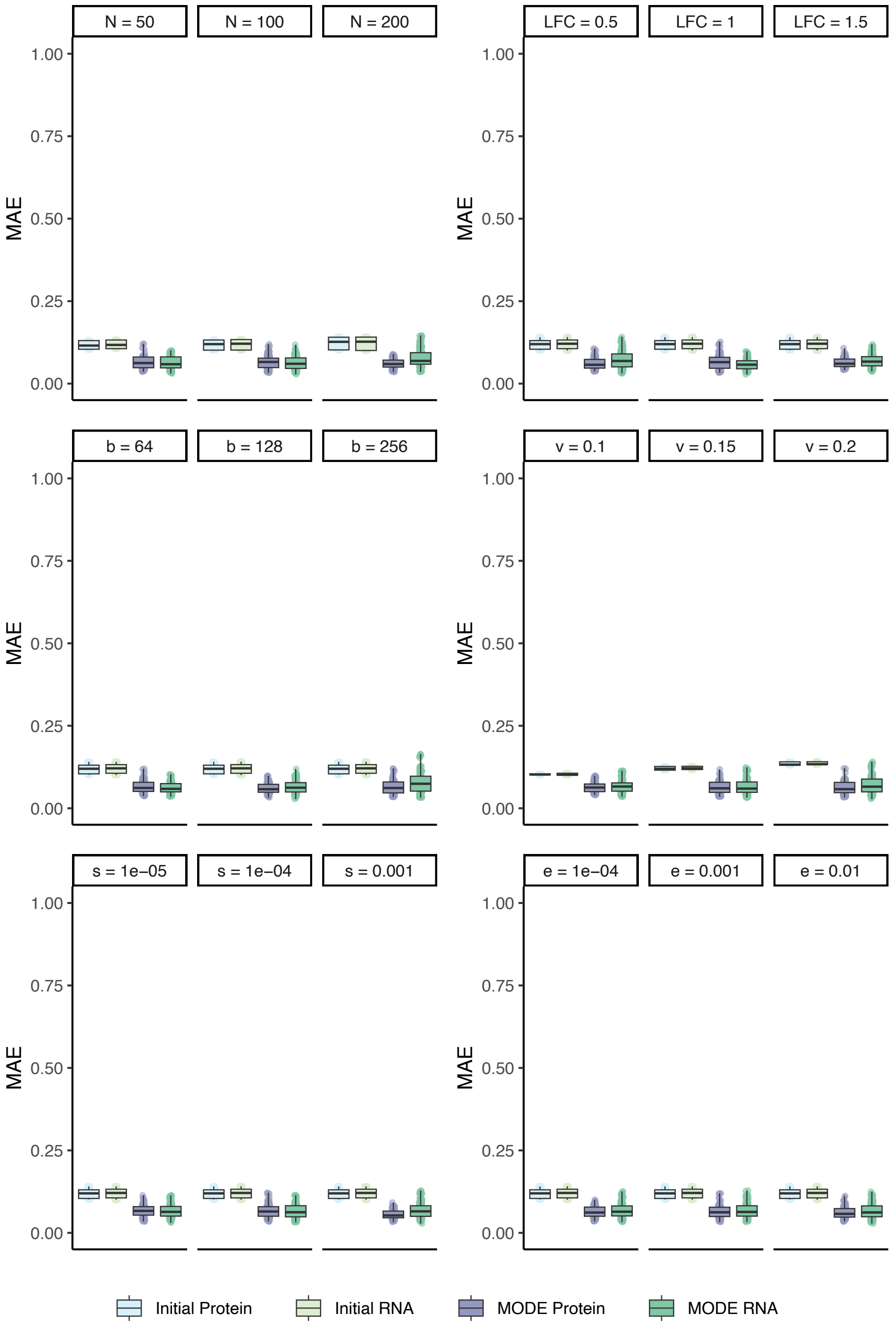


Figure S7: MAE of MODE and the initial cell count fractions in Step 1 JNMF purification by different normal tissue simulation scenarios: sample size (N), log fold change (LFC); and MODE hyperparameters: batch size (b), initial value variance (v), step size (s), and convergence threshold (e).

### 4 Application to pediatric medulloblastoma

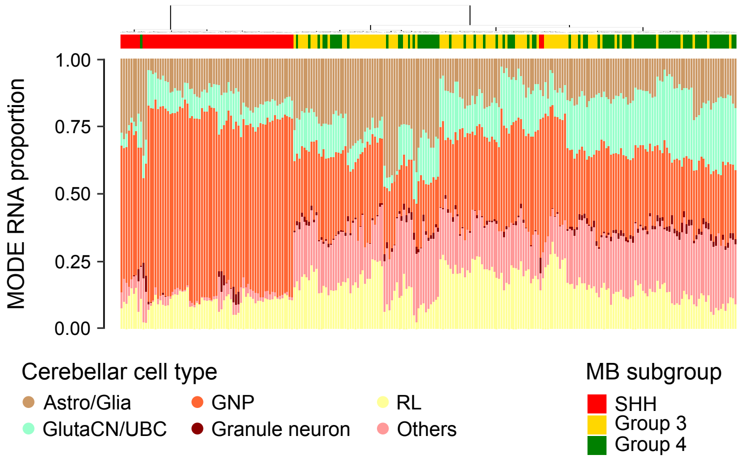


Figure S8: Origin cell proportions in bulk MB transcriptomes resolved by MODE.

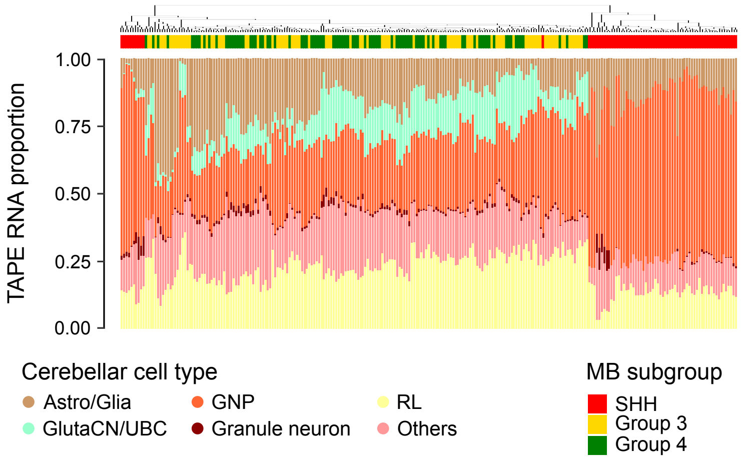


Figure S9: Origin cell proportions in bulk MB transcriptomes resolved by TAPE.

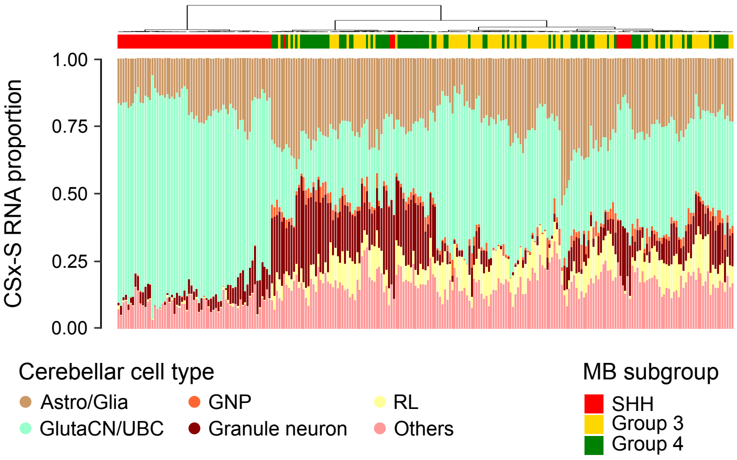


Figure S10: Origin cell proportions in bulk MB transcriptomes resolved by CSx-S.

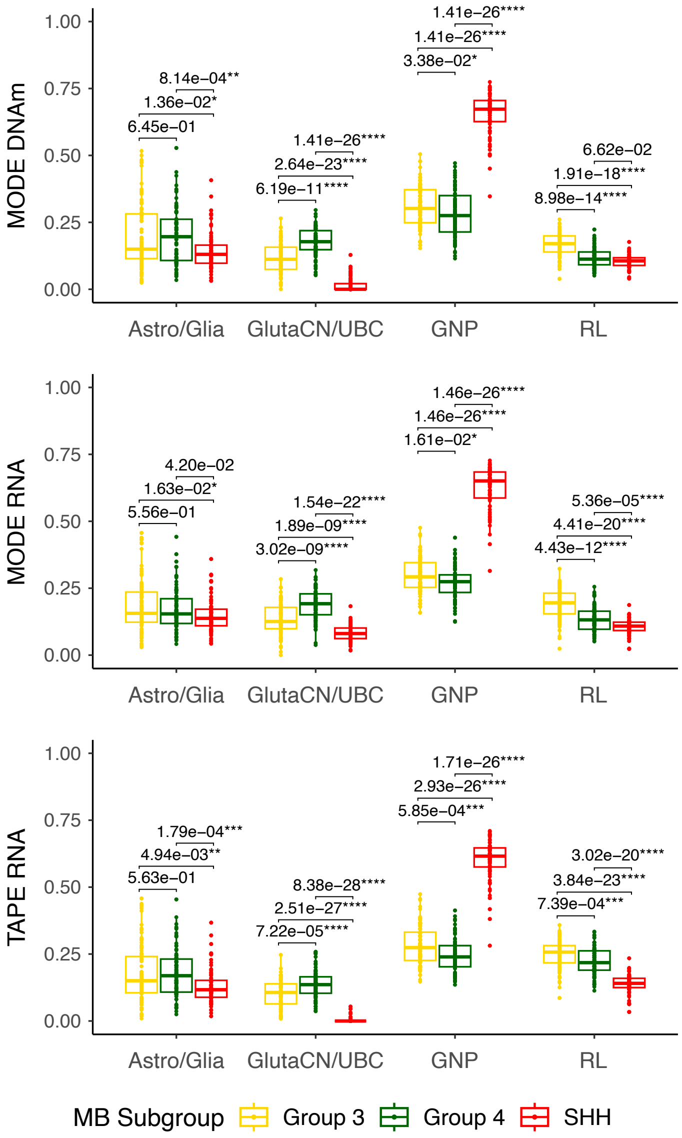


Figure S11: Comparison of MODE (DNAm, RNA) and TAPE (RNA) cell type proportions of Astro/Glia, GlutaCN/UBC, GNP, and RL in MB subgroups.

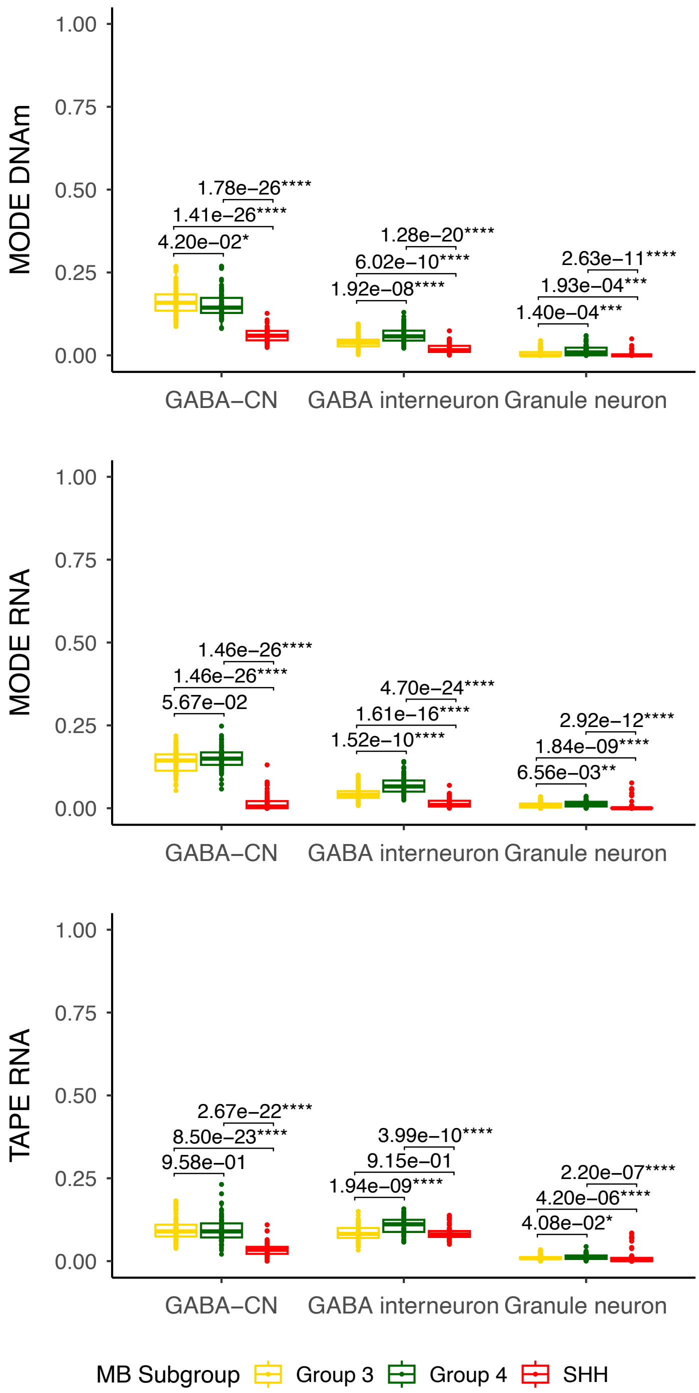


Figure S12: Comparison of MODE (DNAm, RNA) and TAPE (RNA) cell type proportions of GABA-CN, GABA interneuron, and Granule neuron in MB subgroups.

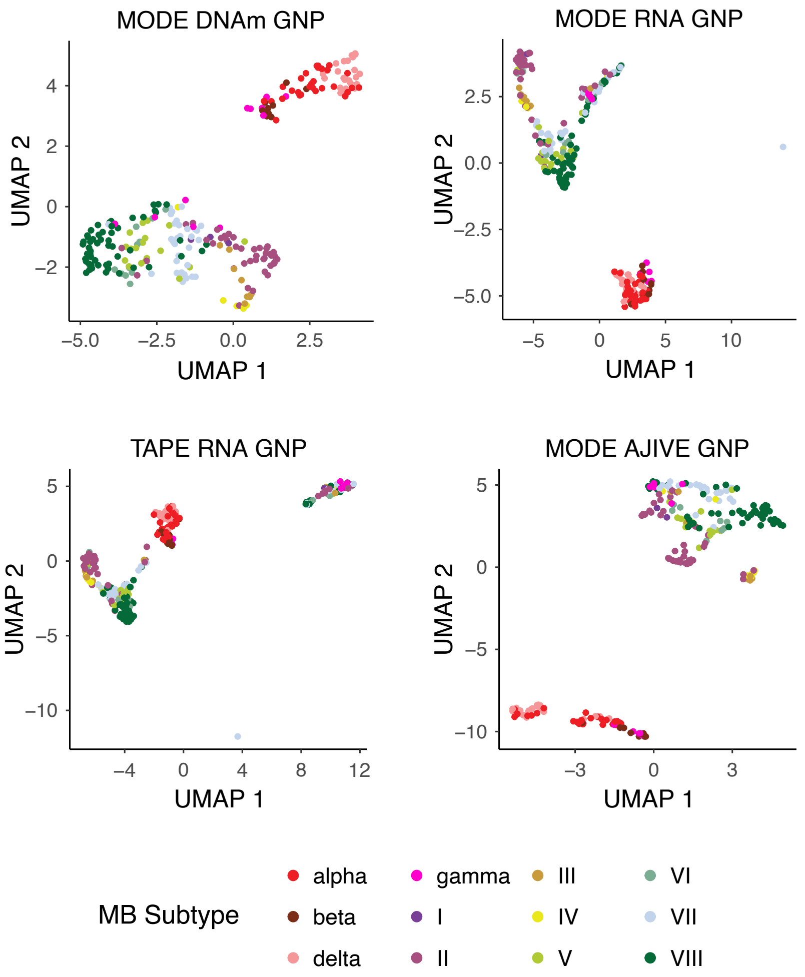


Figure S13: UMAP of MODE (DNAm, RNA), MODE multiomes by AJIVE, and TAPE (RNA) purified profiles mapped to GNP cell state.

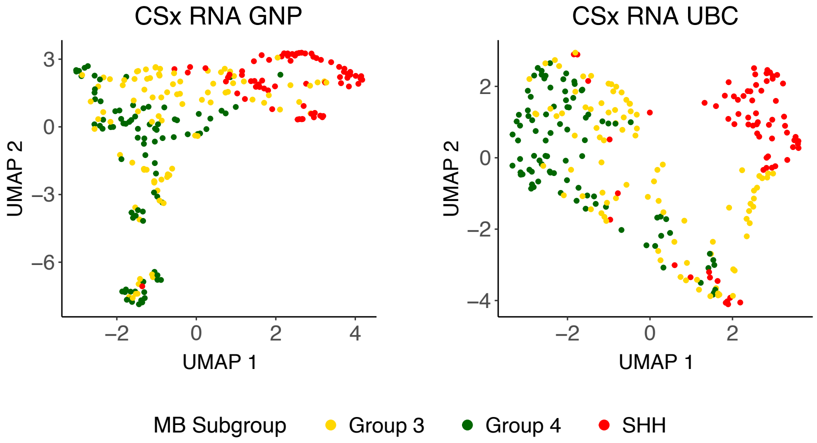


Figure S14: UMAP of CSx RNA purified profiles mapped to GNP, UBC cell state.

### 5 TCGA glioblastoma

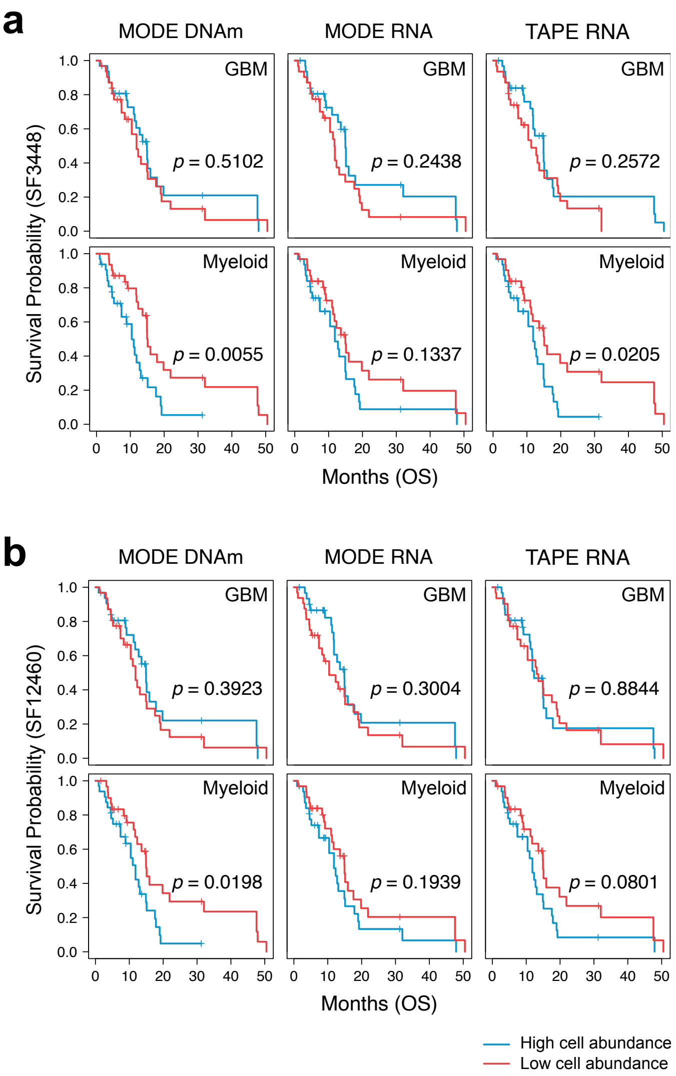


Figure S15: **TCGA GBM patients' overall survival outcomes stratified by binarized cell proportions.** **(a)** Patient sample SF3448 serves as single cell reference. **(b)** Patient sample SF12460 serves as single cell reference.

| | | Event Free Survival | | | | Overall Survival | | | |
|---|---|---|---|---|---|---|---|---|---|
| | | GBM | Myeloid | Neuron | T cell | GBM | Myeloid | Neuron | T cell |
| SF3448 | MODE DNAm | **0.0447** | **0.0032** | 0.6253 | 0.1359 | 0.5102 | **0.0055** | 0.2124 | 0.4130 |
| | MODE RNA | **0.0092** | **0.0388** | 0.4257 | 0.6643 | 0.2438 | 0.1337 | 0.6881 | 0.8490 |
| | TAPE RNA | **0.0291** | **0.0021** | 0.9960 | 0.1367 | 0.2572 | **0.0205** | 0.8548 | 0.3819 |
| SF12460 | MODE DNAm | **0.0277** | **0.0019** | **0.0222** | **0.0251** | 0.3923 | **0.0198** | **0.0115** | 0.0944 |
| | MODE RNA | 0.1682 | **0.0192** | 0.0968 | 0.9565 | 0.3004 | 0.1939 | 0.7678 | 0.8091 |
| | TAPE RNA | 0.4043 | **0.0045** | 0.0801 | 0.1676 | 0.8844 | 0.0801 | 0.1857 | 0.4293 |

Table S1: Log-rank $p$-value of TCGA GBM patients' event free and overall survival stratified by binarized cell proportions in GBM, Myeloid, Neuron, and T cell.

### 6 TCGA breast cancer

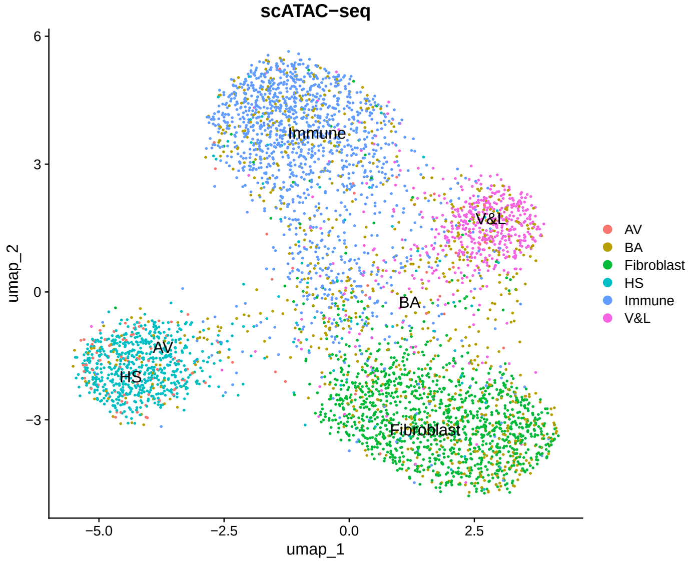


Figure S16: **Human breast cancer scATAC-seq with original cell annotation determined by scRNA-seq.**

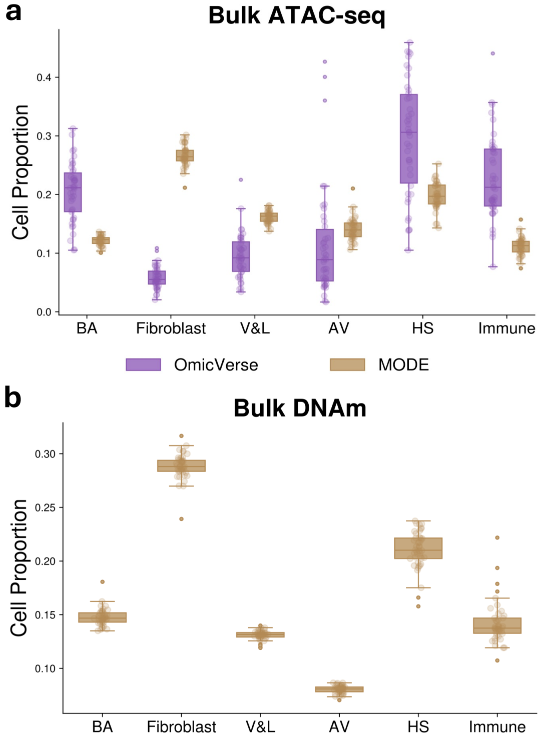


Figure S17: **TCGA breast cancer TME composition using scATAC-seq reference with original cell annotation.** **(a)** TME composition in bulk ATAC-seq resolved by OmicVerse and MODE. **(b)** TME composition in bulk DNAm resolved by MODE.

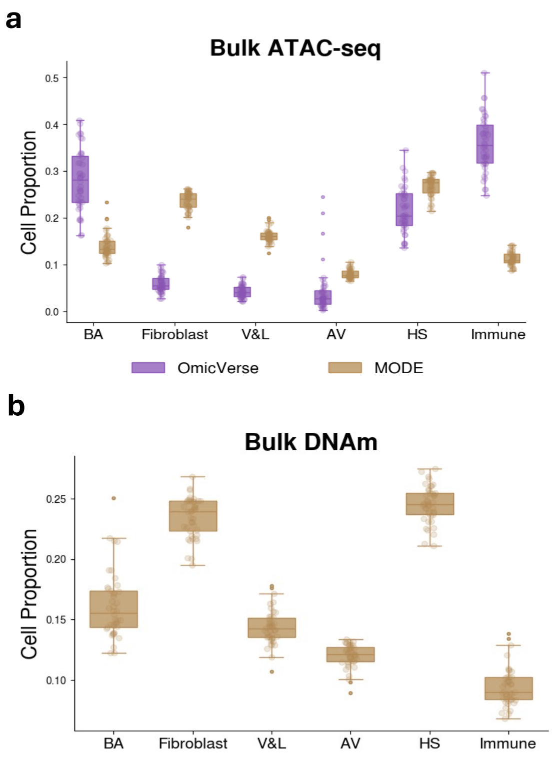


Figure S18: **TCGA breast cancer TME composition using scATAC-seq reference with reannotation.** **(a)** TME composition in bulk ATAC-seq resolved by OmicVerse and MODE. **(b)** TME composition in bulk DNAm resolved by MODE.
